# Cytoplasmic capping enzyme targeted, hypoxia-responsive RNAs, *RORA* and *KCTD16* modulate the aggressiveness of CoCl_2_-induced hypoxic osteosarcoma cells

**DOI:** 10.64898/2026.03.30.715387

**Authors:** Safirul Islam, Utpal Bakshi, Chandrama Mukherjee

## Abstract

Hypoxia is a defining feature of the solid tumour microenvironment and a major determinant of therapeutic response. Hypoxia-inducible factors (HIFs) are central regulators of transcriptional reprogramming under hypoxic stress. Hypoxia can paradoxically elicit both tumour-promoting and tumour-suppressive outcomes, suggesting regulatory mechanisms beyond canonical HIF-dependent pathways. Emerging evidence indicates that hypoxia-responsive RNAs (HRRs) may also be regulated independently of HIFs, with posttranscriptional stabilization playing a critical determinant of hypoxic adaptation. Cytoplasmic mRNA recapping mediated by the cytoplasmic capping enzyme (cCE) has recently emerged as an important post-transcriptional regulatory process, yet its role in hypoxia-driven RNA regulation remains poorly understood. Here, we aimed to identify novel HRRs that modulate cellular adaptability to hypoxia and to determine whether these transcripts are regulated by cCE. Using CoCl_2_-induced hypoxia, we observed a significant reduction in osteosarcoma cell aggressiveness, characterized by decreased proliferation, clonogenic survival, and migratory capacity. Transcriptomic profiling of hypoxic osteosarcoma cells identified *RORA* and *KCTD16* as significantly upregulated and function as suppressors of tumour cell aggressiveness. Integrative *in-silico* CAGE tag analysis followed by cap-specific biochemical assays confirmed that both transcripts are post-transcriptionally stabilized by cCE. Mechanistically, hypoxia-induced stabilization of HIF1α transcriptionally elevated *RORA* and *KCTD16* expression, while cCE further reinforced their stability post-transcriptionally. Stabilization of these cCE-targeted HRRs resulted in suppression of the oncogenic proliferation driver c-Myc, thereby attenuating the aggressive phenotype of hypoxic osteosarcoma cells. Collectively, our findings identify cCE as a previously unrecognized post-transcriptional regulator in hypoxia biology and reveal a RNA-centric mechanism by which hypoxia can restrain tumour aggressiveness.

## INTRODUCTION

Hypoxia is an environmental stress characterized by non-physiological low oxygen levels, typically ≤1% O_2_. In 1955, *Thomlinson* first reported the presence of a distinctly hypoxic core in many malignant tumors (Thomlinson 1977). Subsequent studies by multiple groups established the link between hypoxia and solid tumors, firmly pointing towards hypoxia as a critical driver of tumor progression and cancer pathology.

The 2012 global cancer statistics highlighted the significant mortality burden of cancer, reporting that nearly 20% of all deaths in the United States were cancer-related, with major risk factors including obesity, physical inactivity, and smoking(Torre et al. 2015). Growing evidence indicates that tumor hypoxia promotes cancer progression by activating multiple cellular signaling pathways. Conversely, some studies report that hypoxia can suppress cellular aggressiveness (Gardner et al. 2001; Kumar and Vaidya 2016; Shi et al. 2023). This duality makes the role of hypoxia in cancer paradoxical and an active area of investigation.

Cellular adaptation to hypoxia is primarily governed by the oxygen-sensing transcription factor HIF1α, which reprograms the transcriptome to facilitate their endurance under low-oxygen conditions. This reprogramming involves diverse hypoxia-responsive RNAs (HRRs), including both coding and noncoding transcripts (Parodi et al. 2018; Abou Khouzam et al. 2023; Reheman et al. 2024). In addition to HIF1α, several noncoding RNAs such as microRNAs (miRNAs), long noncoding RNAs (lncRNAs), and circular RNAs (circRNAs), along with epigenetic factors, also regulate cellular responses to hypoxia (Islam and Mukherjee 2022). Since the effective hypoxic adaptation depends on these HRRs, their stability is a key determinant of cellular responses during hypoxic stress.

Among the various posttranscriptional modifications that ensure the stability of RNA Pol II-derived transcripts, 5′-capping is particularly critical, as it protects RNA 5′ ends from exonucleolytic degradation and facilitates processes such as splicing and 3′-polyadenylation (Inoue et al. 1989; Lewis et al. 1996; Flaherty et al. 1997; Shuman 2002). Although initially thought to occur exclusively in the nucleus, the capping enzyme RNGTT (CE) is now known to also function in the cytoplasm. In this context, CE in concert with RNMT, RAMAC, 5′-RNA kinase and the adaptor protein Nck1 to reconstitute cap structures on decapped RNAs, thereby restoring their stability and translational competence (Mukherjee et al. 2012; Mukherjee et al. 2014; Trotman et al. 2017). Importantly, accumulating evidence suggests that transcripts containing internal CAGE tags can serve as substrates for cytoplasmic capping enzyme (cCE) (Kiss et al. 2015). Subsequent studies have identified several endogenous cCE targets, including both mRNAs and a subset of long noncoding RNAs (Del Valle Morales et al. 2020; Mukherjee et al. 2023).

Current evidence indicates that beyond its role in 5′-capping, the capping enzyme (CE/RNGTT) has broader biological functions. CE has been implicated in regulating Hedgehog signalling during *Drosophila* development and in the pathogenesis of certain cancers (Chen et al. 2017; Borden et al. 2021). Notably, in non-small cell lung cancer (NSCLC), CE expression and activity are modulated by the m^6^A methyltransferase METTL3, revealing an additional layer of epitranscriptomic regulation of CE-mediated RNA processing (Del Valle-Morales et al. 2024). Given the nuclear origin of cCE, it was hypothesized that cCE may participate in biological processes beyond canonical mRNA capping. More recently, we demonstrated that CoCl_2_ induced hypoxic stress globally modulates CE expression, impacting both nuclear and cytoplasmic pools. Under hypoxic conditions, several hypoxia-responsive lncRNAs lose their 5′-caps and become destabilized; however, cytoplasmic CE counteracts this effect by re-capping these transcripts, thereby preserving their stability (Islam and Mukherjee 2025). Collectively, these findings underscore a critical role for CE/cCE in regulating cellular responses to hypoxia and highlight its importance in hypoxia-driven transcriptomic adaptation.

Hypoxia can be modelled *in vitro* by physical induction (≤1% O_2_ using hypoxia chambers) or chemical induction with hypoxia mimetics such as CoCl_2_ or deferoxamine (Muñoz-Sánchez and Chánez-Cárdenas 2019). Comparative transcriptomic analyses show that these models elicit both overlapping and distinct gene expression patterns, underscoring the context-dependent nature of hypoxic responses influenced by the mode, duration, and severity of oxygen deprivation (Calvo-Anguiano et al. 2018). Several studies have reported that hypoxia can suppress cellular aggressiveness in models such as multiple myeloma, retinoblastoma, and mouse embryonic stem cells (Muz et al. 2014; Binó et al. 2016; Yang et al. 2017). In line with this, one study demonstrated that hypoxic conditions significantly reduce the tumor-forming potential of osteosarcoma cells, suggesting that low oxygen levels can act as a restraining factor in certain cancer contexts (Zhang et al. 2013). However, the molecular mechanisms behind this suppressive effect remains unclear. Recent research has begun to shed light on potential pathways, highlighting the role of posttranscriptional regulatory processes.

Notably, our previous work connected hypoxia to CE/cCE-mediated post-transcriptional stabilization of lncRNAs, adding a new layer to this regulatory network. Building on these observations and addressing key gaps in hypoxia research, the present study investigates whether hypoxia-responsive cytoplasmic capping enzyme (cCE)-targeted RNAs contribute in regulating osteosarcoma cell aggressiveness under CoCl_2_ induced hypoxia. We demonstrate that CoCl_2_ mediated hypoxic stress significantly suppresses the proliferative, clonogenic, invasive, and metastatic potential of osteosarcoma cells, as validated by multiple functional assays. Hypoxia also induced a pronounced cell-cycle arrest, characterized by a marked reduction in the S-phase population and accumulation of cells in the G2/M phase. Transcriptomic profiling identified *RORA* (Retinoic Acid Receptor-Related Orphan Receptor Alpha) and *KCTD16* (Potassium Channel Tetramerization Domain Containing 16) as a novel hypoxia-responsive transcripts that act as substrates of the cytoplasmic capping enzyme. Functional analyses revealed that both RNAs are essential for the hypoxia-induced suppressive phenotype. Mechanistically, *RORA* and *KCTD16* proteins attenuated c-Myc expression, as confirmed by reciprocal knockdown and overexpression experiments, leading to reduced proliferative capacity and cell-cycle arrest. Consistently, depletion of either transcript enhanced cellular aggressiveness, whereas their ectopic overexpression suppressed it. Collectively, this study provides the first evidence that modulation of cCE-targeted RNA stability can directly influence hypoxia-driven cellular outcomes, uncovering a new post-transcriptional regulatory axis in hypoxic osteosarcoma cells.

## RESULTS

### CoCl_2_ induced hypoxia reduces osteosarcoma cellular aggressive phenotypes

The impact of hypoxia on cellular behaviour has been extensively investigated worldwide, with numerous studies indicating that a hypoxic microenvironment promotes tumorigenesis (Terry et al. 2018; Tang et al. 2021; Kunachowicz et al. 2025), while other reports demonstrate that hypoxia can significantly inhibit cellular aggressiveness (Gardner et al. 2001; Binó et al. 2016; Kumar and Vaidya 2016; Shi et al. 2023). Therefore, these observations prompted us to examine the effect of CoCl_2_ induced hypoxia on osteosarcoma cells.

To examine the effects of CoCl_2_ induced hypoxia on osteosarcoma cells, we employed a previously established hypoxic osteosarcoma model in which U2OS and MG63 osteosarcoma cells were treated with 200 µM CoCl_2_ for 24 hrs (Islam and Mukherjee 2025). To evaluate the impact of CoCl_2_ induced hypoxia on osteosarcoma cell aggressiveness, we assessed key parameters including proliferative capacity, colony-forming ability, migratory behaviour, and invasive potential using the assays described in the Materials and Methods section. In this study, we observed that CoCl_2_ induced hypoxia significantly reduced the proliferative potential of both U2OS and MG63 osteosarcoma cells, as indicated by a marked decrease in BrdU-positive cells in hypoxic conditions compared with control cells (Fig. 1A-D). These findings were further supported by clonogenic assays, which demonstrated that the reduced proliferative capacity of hypoxic osteosarcoma cells was accompanied by a significant impairment in their colony-forming ability (Fig. 1E-H). Moreover, CoCl_2_ induced hypoxia significantly reduced the migratory capacity of both U2OS and MG63 cells compared with untreated controls (Fig. 1I–L). Importantly, our data demonstrated that hypoxic U2OS cells exhibited a reduced invasive potential, as evidenced by the Boyden chamber invasion assay (Supplemental Fig. 1A-B). These findings led us to examine cell cycle dynamics under the same conditions. Flow cytometry based analysis using propidium iodide revealed that CoCl_2_ induced hypoxia reduced the proportion of cells in S phase (DNA synthesis) and led to an accumulation of cells in the G2/M phase in osteosarcoma cells (Supplemental Fig. 1C-F). Since BrdU incorporation occurs exclusively in actively proliferating S-phase cells, the reduced number of BrdU-positive cells directly corresponds with the observed decrease in the S-phase population in the cell-cycle analysis. Likewise, the hypoxia-induced G2/M arrest is consistent with the diminished clonogenic capacity of osteosarcoma cells under these conditions. Collectively, these results demonstrate that exposure to 200 µM CoCl_2_ for 24 hrs markedly suppresses the aggressive phenotypes of both U2OS and MG63.

**FIGURE 1.**
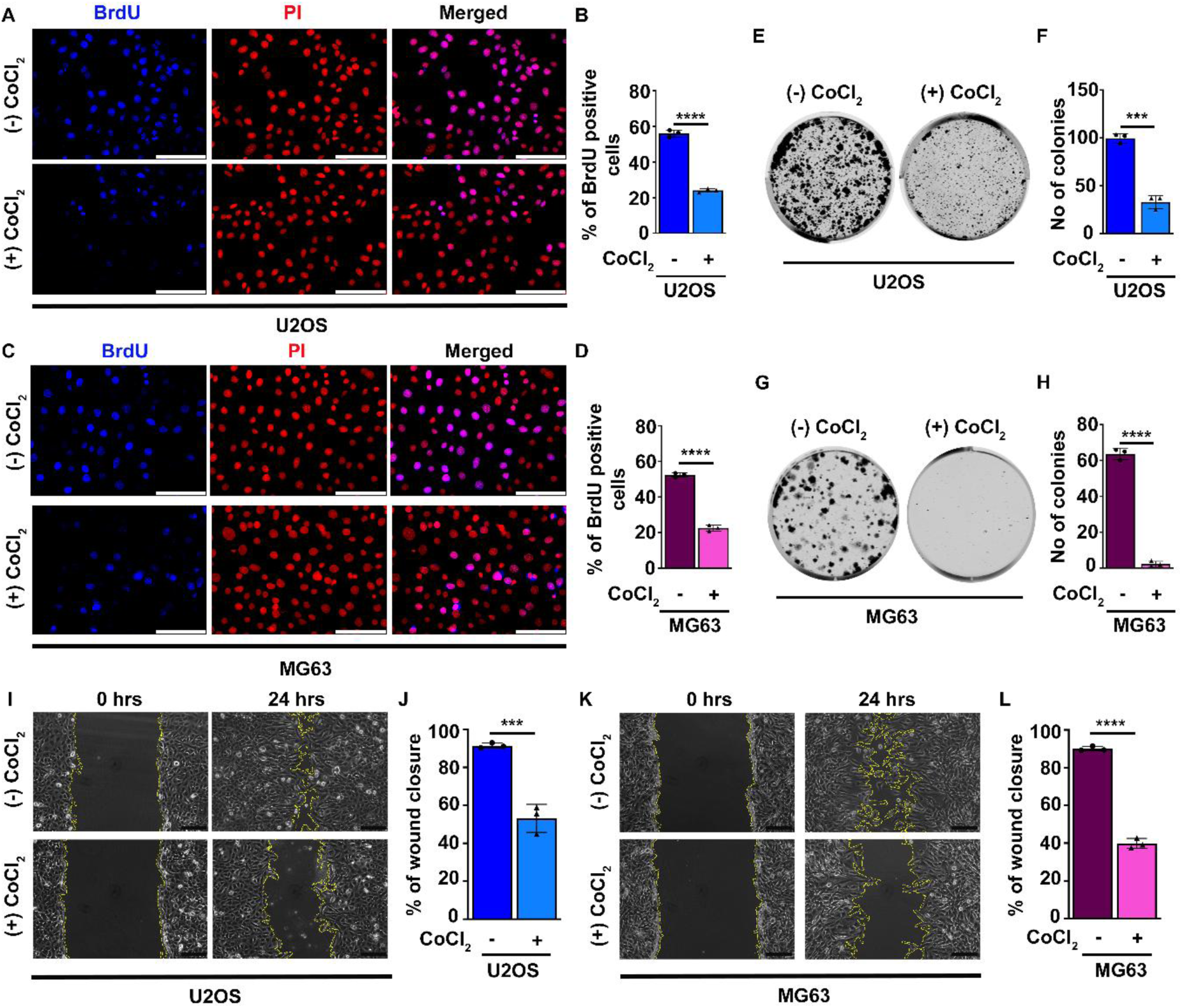
CoCl_2_ induced hypoxia compromises the cellular aggressiveness of osteosarcoma cells. (*A* and *C*) Representative microscopic images of BrdU incorporation assays in U2OS and MG63 osteosarcoma cells under normoxic (control) and CoCl_2_ induced hypoxic conditions. Scale bar = 50 μm. (*B* and *D*) Quantification of BrdU-positive cells showing a significant decrease in proliferation in hypoxic U2OS and MG63 cells compared with their respective controls (n ≥ 20 cells per condition). (*E* and *G*) Representative images of colony-formation assays in control and CoCl₂-treated hypoxic U2OS and MG63 cells, respectively. (F and H) Quantification of colony numbers showing a marked reduction in the clonogenic potential of hypoxic osteosarcoma cells. (*I* and *K*). Representative images of cell-migration assays in U2OS and MG63 cells under control and hypoxic conditions. Scale bar = 50 µm. (*J* and *L)* Quantification of migrated cells showed a significant reduction in the migratory capacity of hypoxic U2OS and MG63 cells, respectively. Statistical significance was calculated using two-tailed Student’s *t*-test and is represented as mean ± SD from three biological replicates. ns: non-significant, *P < 0.05, **P < 0.005, ***P < 0.0005, ****P < 0.0001.

### *RORA* and *KCTD16* are potential regulators of the suppressed aggressive phenotypes observed in CoCl_2_ induced hypoxic U2OS cells

Previously, we observed that CoCl_2_ induced hypoxia significantly reduces the aggressiveness of osteosarcoma cells. A similar finding was reported by Zhang et al. (2013), who showed that CoCl_2_ markedly diminishes the tumor-forming potential of osteosarcoma cells both *in vitro* and *in vivo* (Besnard et al. 2002; Chauvet et al. 2002; Chauvet et al. 2004; Zhang et al. 2013). However, their study was limited to phenotypic observations and did not explore the underlying mechanisms. Building on these observations and our own data, we aimed to identify the key molecular players mediating the suppressive oncogenic phenotypes imparted by CoCl_2_ induced hypoxia in osteosarcoma cells *in vitro*.

To identify the molecular drivers of CoCl_2_ induced hypoxic responses in osteosarcoma cells, we performed whole-transcriptome analysis (GSE293541) using cytoplasmic RNA from control and CoCl_2_ treated U2OS cells. Among the differentially expressed genes, highlighted in the heat map (Fig 2A), functional enrichment analysis (Fig 2B) and volcano plot (Fig 2C), we focused on *RORA* (RAR Related Orphan Receptor A) and *KCTD16* (Potassium Channel Tetramerization Domain Containing 16) due to their upregulation under hypoxia and relevance to hypoxia-associated pathways (Shi et al. 2022; Zhang et al. 2025). Importantly, TCGA pan-cancer data show that both *RORA* and *KCTD16* are broadly downregulated across multiple tumor types (Supplementary Fig 2 A, B). *RORA* has been reported to suppress proliferation in cancers such as glioma, endometrial cancer, and esophageal squamous cell carcinoma (Sun et al. 2018; Jiang et al. 2020; Ma et al. 2021), but its function in hypoxic osteosarcoma remains unclear. In contrast, the role of *KCTD16* in cancer or hypoxia has not been explored however the other family member with similar structure that is *KCTD2* has been known to inhibit gliomagenesis (Kim et al. 2017). As a positive control for hypoxia-related responses, *BNIP3* (BCL2 Interacting Protein 3) was included throughout the experiments due to its well-established association with hypoxia signalling (Noordeen and Lakshmi 2026). HIF1α is the master regulator of hypoxia; therefore, the presence of HIF1α or its downstream targets (e.g., *EPO*, *CA9*, *VEGFA*) is widely used as a hypoxic marker (Calvo-Anguiano et al. 2018). To validate the CoCl_2_ induced hypoxic osteosarcoma model, *CA9* expression was examined at the transcript level and HIF1α at the protein level (Fig 2D-F). The RNA-seq findings were further validated by qPCR and western blot analyses, which showed increased expression of the selected transcripts in CoCl_2_ treated hypoxic U2OS cells, in strong agreement with the RNA-seq data (Fig 2D-F). Finally, these results strongly support that CoCl_2_ induced hypoxia induces RORA and KCTD16 expression.

**FIGURE 2.**
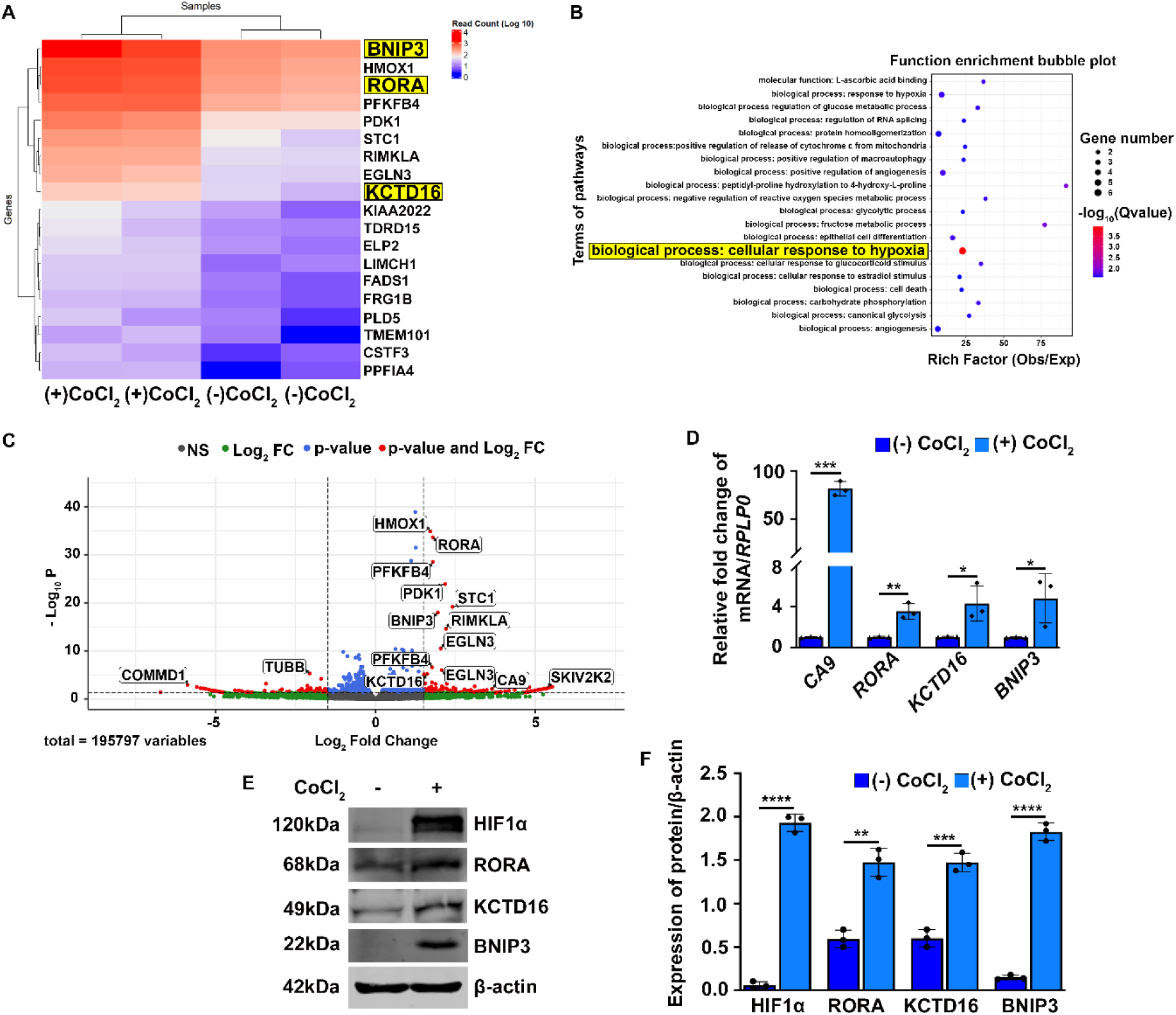
Transcriptomic profiling of CoCl_2_ treated hypoxic U2OS osteosarcoma cells. (*A*) Heat map showing differentially expressed genes, (*B*) Functional enrichment bubble plot highlighting hypoxia-associated biological pathways, (*C*) Volcano plot depicting significantly upregulated (red) and downregulated (blue) genes (p ≤ 0.05, log2FC ≥ 1). (D) qPCR validation of selected cytoplasmic transcripts, corroborating RNA-seq results; *CA9* served as a positive hypoxia marker. (*E*) Western blot analysis of selected proteins under hypoxia. (*F*) Densitometric quantification of immunoblots using ImageJ, consistent with qPCR data; HIF1α protein confirms hypoxia induction, and β-actin was used for normalization. Statistical significance was calculated using two-tailed Student’s *t*-test and is represented as mean ± SD from three biological replicates. ns: non-significant, *P < 0.05, **P < 0.005, ***P < 0.0005, ****P < 0.0001.

### *RORA* and *KCTD16* contain internal CAGE tags and posttranscriptionally regulated by cCE

Previous studies have shown that several mRNAs are posttranscriptionally stabilized by cCE (Mukherjee et al. 2012). More recently, we demonstrated that CoCl_2_ induced hypoxia globally increases CE expression, including both nuclear CE and cytoplasmic CE, and that the elevated cCE selectively stabilizes a subset of hypoxia-responsive lncRNAs during hypoxic stress(Islam and Mukherjee 2025). These observations suggest that CE/cCE has broader, still incompletely understood roles in hypoxia, prompting us to investigate whether the hypoxia-responsive transcripts *BNIP3*, *RORA*, and *KCTD16* are also cCE targets. To test whether the selected transcripts *BNIP3*, *RORA*, and *KCTD16* are substrates of cCE, we assessed their steady-state expression upon cCE inhibition. Direct knockdown of cCE would disrupt essential nuclear CE activity; therefore, we used a U2OS cell line stably expressing a doxycycline-inducible dominant-negative cytoplasmic capping enzyme mutant (K294A) (Islam and Mukherjee 2025). Nucleo-cytoplasmic fractionation was validated by immunoblotting for Lamin A/C (nuclear) and GAPDH (cytoplasmic), confirming clean separation. Myc-K294A expression was detected only in the cytoplasm upon doxycycline induction (Fig 3A). qPCR and immunoblot analysis of cytoplasmic RNA and protein revealed that RORA and KCTD16 levels were significantly reduced upon cCE inhibition (Fig 3 B-D), whereas BNIP3 remained unchanged (Fig 3 B), indicating that *RORA* and *KCTD16* are specifically stabilized by cCE. These results indicate that *RORA* and *KCTD16* rely on cCE for posttranscriptional stability, as cCE inhibition markedly reduces their cytoplasmic levels, whereas *BNIP3* appears unaffected and is likely not a cCE substrate.

**FIGURE 3.**
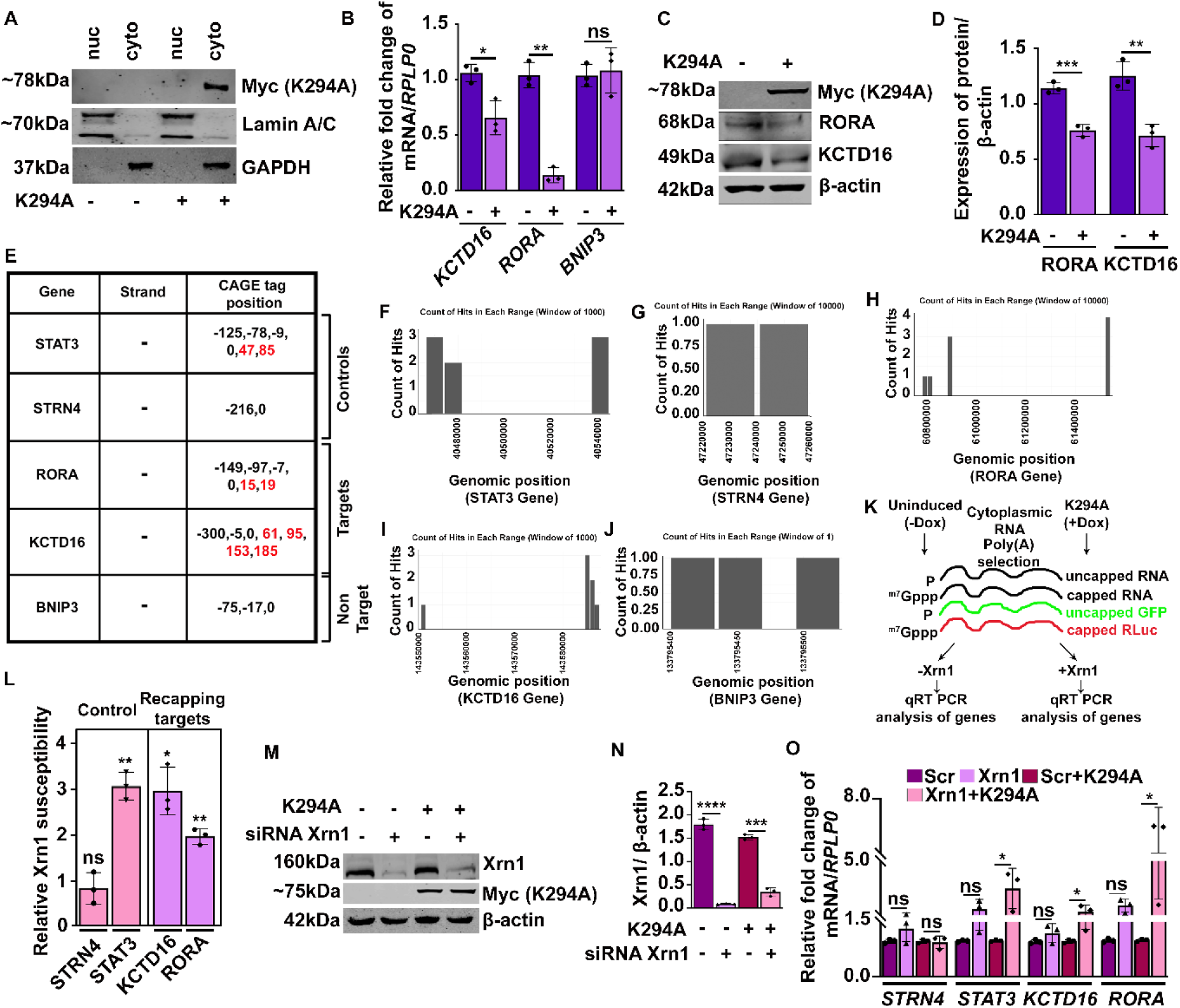
*RORA* and *KCTD16* are posttranscriptionally regulated by cCE. (*A*) U2OS cells, either stably expressing K294A upon doxycycline induction or uninduced controls, were biochemically fractionated into nuclear and cytoplasmic compartments. Western blot analysis confirmed fractionation quality using Lamin A/C as a nuclear marker and GAPDH as a cytoplasmic marker. Myc blotting verified the expression of K294A upon doxycycline induction. (*B*) Quantitative real-time PCR analysis revealed a significant reduction in the cytoplasmic levels of *RORA* and *KCTD16* transcripts in K294A expressing cells compared with controls, while *BNIP3* levels remained unchanged. (*C*) Western blot analysis further confirmed decreased protein levels of RORA and KCTD16 in K294A-expressing cells. Myc blotting verified stable K294A expression. (*D*) Densitometric analysis of RORA and KCTD16 protein bands from panel C was performed using ImageJ software. β-Actin was used as a loading control for normalization. Statistical analysis was calculated by performing two-tailed Student’s *t*-test. (*E*) Table summarizing the internal CAGE (Cap Analysis of Gene Expression) sites identified within the analysed transcripts, with the specific positions highlighted in red. (*F*-*J*) Bar graphs representing the genomic distribution of CAGE peaks for each gene, illustrating the relative frequency of CAGE signals across different transcript regions. (*K*) Schematic illustration of the Xrn1 susceptibility assay used to assess the stability of 5′-capped transcripts. (*L*) Relative 5′-end loss of *RORA* and *KCTD16* was assessed using an in vitro Xrn1 susceptibility assay. In K294A-expressing cells, both transcripts exhibited a level of 5′-end loss comparable to *STAT3*, a known cCE target, relative to control cells. Statistical analysis was performed using one sample Student’s *t*-test. (*M*) Western blot analysis showing Xrn1 protein levels in Xrn1 knockdown cells with or without doxycycline-induced K294A expression. Myc detection confirmed successful induction of the dominant-negative cCE mutant. (*N*) Quantification of Xrn1 knockdown efficiency was performed using ImageJ software, with β-Actin serving as the internal loading control. (*O*) *RORA* and *KCTD16* exhibited the most pronounced rescue in Xrn1 knockdown cells expressing K294A, indicating their strong dependence on cytoplasmic capping for stability. Statistical significance was determined using one-way ANOVA. All the data is represented as mean ± SD from three biological replicates. ns: non-significant, *P < 0.05, **P < 0.005, ***P < 0.0005, ****P < 0.0001.

Next, we investigated if *RORA* and *KCTD16* contain internal CAGE tags that was used before to determine if the transcripts undergo cytoplasmic capping (Kiss et al. 2015; Berger et al. 2019; Mukherjee et al. 2023). I*n silico* analysis for internal CAGE tags, using *STAT3* as a positive control and *STRN4* as a non-target (Mukherjee et al. 2012), revealed that *RORA* and *KCTD16* contain internal CAGE tags downstream of their canonical TSS (transcription start site), similar to *STAT3*. In contrast, *BNIP3* and *STRN4* lacked such internal tags. The genomic coordinates and distribution of these CAGE sites (Fig 3 E-J) support the idea that *RORA* and *KCTD16* are likely bona fide substrates of the cCE complex. Therefore, we speculated that the reduced abundance of these transcripts in K294A-expressing cells results from the loss of their 5′ caps under cytoplasmic capping-inhibited conditions. In the absence of active cCE, these RNAs fail to undergo cytoplasmic recapping and consequently become susceptible to Xrn1-mediated 5′-to-3′ exonucleolytic degradation.

To test this hypothesis, we performed an Xrn1 susceptibility assay, which differentiates capped from uncapped transcripts (Mukherjee et al. 2012; Mukherjee et al. 2023; Gayen et al. 2024; Islam and Mukherjee 2025). Poly(A)-selected cytoplasmic RNAs from control and K294A-expressing cells were spiked with capped Renilla luciferase RNA and synthetic uncapped GFP RNA, followed by incubation in the presence or absence of Xrn1. Subsequent qPCR was performed using gene-specific primers, as outlined in Fig 3 K. Ct values were normalized to the capped Renilla control which remained same throughout the tratment. Xrn1 susceptibility was calculated as the ΔX_K294A_/ΔX_Control_ ratio, where ΔX_Control_ represents the change in Xrn1 sensitivity in control cells and ΔX_K294A_ represents the corresponding change in K294A-expressing cells as done previously (Gayen et al. 2024; Islam and Mukherjee 2025). Strikingly, *RORA* and *KCTD16* displayed a pronounced increase in Xrn1 susceptibility, comparable to the established cCE target *STAT3* (Fig 3 L). In contrast, the non-target *STRN4* showed no change, confirming the specificity of the assay (Fig 3 L). These results demonstrate that inhibition of cytoplasmic capping leads to the accumulation of uncapped *RORA* and *KCTD16* transcripts, rendering them vulnerable to Xrn1-mediated degradation. Under normal conditions, active cCE preserves the stability of these transcripts by restoring their 5′ caps.

To further corroborate the Xrn1 susceptibility results, we hypothesized that the reduced levels of *RORA* and *KCTD16* observed in doxycycline-induced K294A-expressing cells (Fig 3 C-D) result from Xrn1-mediated degradation of uncapped transcripts. Accordingly, depletion of Xrn1 should restore their expression under cCE-inhibited conditions. To test this, K294A cells were transfected with either scrambled siRNA or siXrn1, and transcript levels were assessed. Western blot analysis confirmed efficient Xrn1 knockdown, with more than 85% reduction in protein levels (Fig 3 M and N). Consistent with our hypothesis, qPCR analysis revealed a significant rescue of *RORA* and *KCTD16* expression in the cytoplasmic fraction of Xrn1-depleted K294A cells, comparable to the established cCE target *STAT3* (Fig 3 O). In contrast, *STRN4*, a known cCE non-target, showed no change, underscoring the specificity of the effect (Mukherjee et al. 2012).

Importantly, we observed that the steady state expression of *RORA* and *KCTD16* was markedly reduced in K294A cells treated with CoCl_2_ compared to untreated K294A cells (Supplemental Fig. 2C). This suggest that the decreased expression of these trasnscripts under hypoxic condition in K294A cells may result from to the loss of 5’ cap rendering them susceptible to Xrn1-mediated 5′-3′ exoribonucleolytic degradation. Consequently, their transcript levels are significantly compromised in hypoxic K294A cells. Notably, this observation is consistent with our previous findings, where we demonstrated that hypoxic stress induces cap loss in a subset of hypoxia-responsive lncRNAs; however, their sustained expression under hypoxia is maintained through cCE-mediated recapping (Islam and Mukherjee 2025).

Collectively, these results demonstrate that inhibition of cytoplasmic capping leads to the accumulation of uncapped *RORA* and *KCTD16* transcripts, which are subsequently degraded by the 5′-3′ exonuclease Xrn1. In contrast, under conditions where cCE is active, these transcripts are efficiently recapped, thereby protected from exonucleolytic decay. Thus, *RORA* and *KCTD16* are established as bona fide cCE-dependent RNA targets whose stability is maintained through cytoplasmic recapping.

### Suppression of HIF1α down regulates RORA and KCTD16 expression in osteosarcoma cells

RNA-seq analysis and its validation confirmed that CoCl_2_ induced hypoxia significantly increased RORA and KCTD16 expression in osteosarcoma cells. Previously we have shown that HIF1α during CoCl_2_ induced hypoxia elevates expression of cCE which also stabilizes the target transcripts (Islam and Mukherjee 2025). We wonder if hypoxia-mediated upregulation of *RORA* and *KCTD16* were regulated by HIF1α and cCE protects these transcripts during hypoxic conditions.

To investigate the role of HIF1α in regulating the expression of *RORA* and *KCTD16*, HIF1α expression and activity were inhibited using PX478, a well-established pharmacological inhibitor of HIF1α (Islam and Mukherjee 2025). PX478 suppresses HIF1α at both transcriptional and translational levels and further destabilizes the protein by enhancing its ubiquitination while inhibiting deubiquitination (Koh et al. 2008). U2OS cells were treated with PX478 at a final concentration of 40 µM, either in the presence or absence of 200 µM CoCl_2_, for 24 hrs. Changes in the expression of the validated candidate genes were subsequently assessed at both the mRNA and protein levels using qPCR and western blotting, respectively.

As expected, *BNIP3*, the known target of HIF1α (Noordeen and Lakshmi 2026) showed reduced expression in transcript as well as protein levels when treated with PX478 in addition to CoCl_2_ as compared to the respective U2OS control cells (Fig 4, A, C, and D). Similarly, a significant reduction in the expression of RORA and KCTD16 was found at both RNA and protein levels in the PX478-treated hypoxic group as compared to the control (Fig 4.3, A, C, and D). The efficacy of PX478 was assessed by measuring *CA9* transcript levels and HIF1α protein expression using qPCR and western blotting, respectively. Cells treated with both CoCl_2_ and PX478 exhibited markedly reduced HIF1α protein levels compared with cells treated with CoCl_2_ alone (Fig 4C and D). A similar trend was observed at the transcript level, with decreased *CA9* expression in CoCl_2_ + PX478-treated cell lysates (Fig 4A). These findings were further validated in another osteosarcoma cell line, MG63, which displayed responses comparable to those observed in U2OS cells (Fig 4B, E, and F). Collectively, these data demonstrate that the expression of RORA and KCTD16 is driven by HIF1α during CoCl_2_ induced hypoxic adaptation in osteosarcoma cells.

**FIGURE 4.**
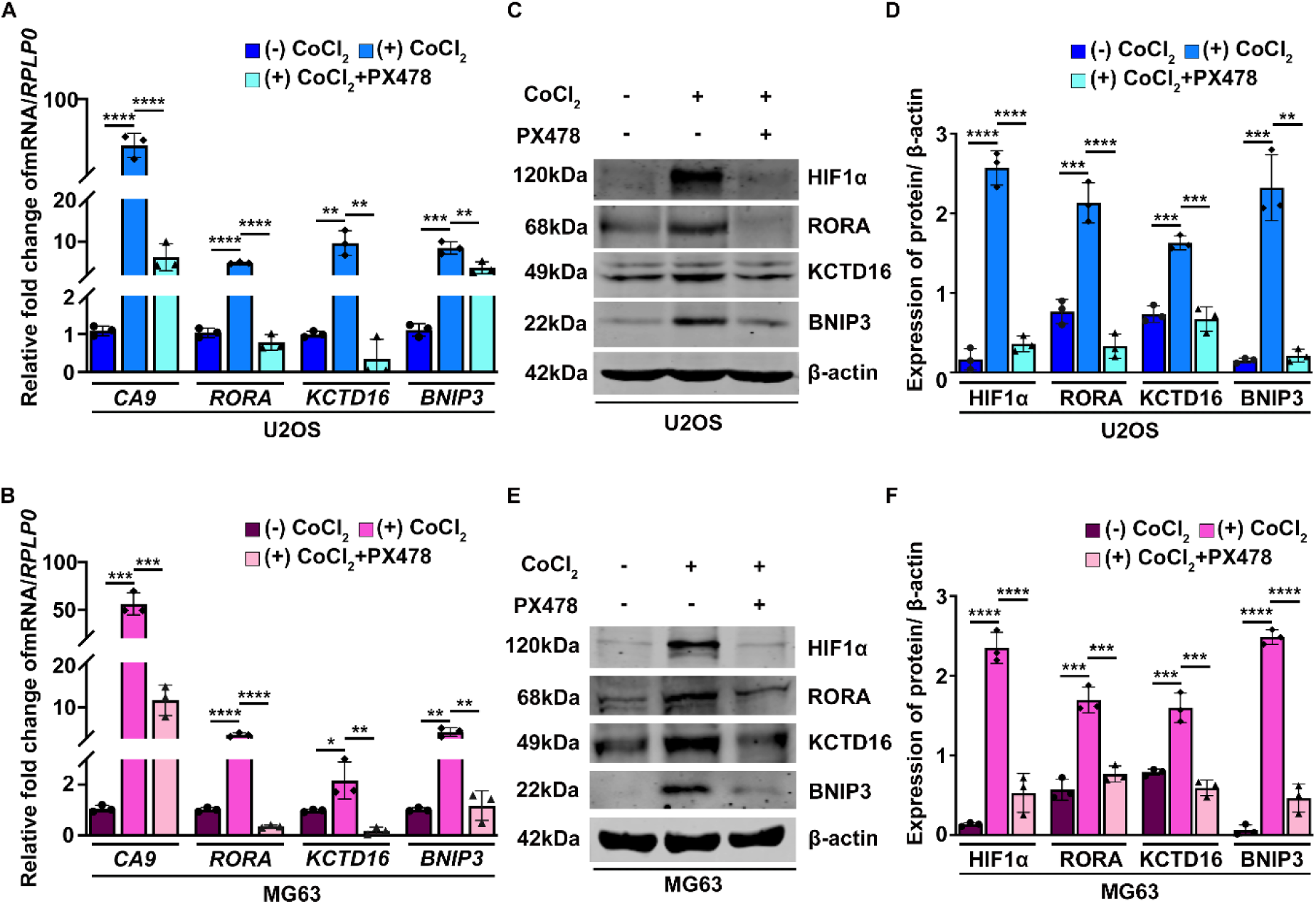
HIF1α regulates the expression of the selected transcripts under CoCl_2_ induced hypoxic conditions. (*A* and *B*) qPCR analysis of the selected transcripts in U2OS and MG63 cells revealed reduced expression following treatment with the HIF1α inhibitor PX478, irrespective of hypoxia induction. (*C* and *E*) Western blot analysis of U2OS and MG63 cells demonstrated reduced protein levels of all selected targets, including HIF1α, in PX478-treated hypoxic samples. (*D* and *F*) Quantification of western blot band intensities corresponding to panels C and E was performed using ImageJ software. β-Actin served as the loading control for normalization. Data are presented as mean ± SD from three biological replicates. Statistical significance was determined using one-way ANOVA. ns, not significant; *P < 0.05; **P < 0.005; ***P < 0.0005; ****P < 0.0001.

### HIF1α is recruited to the hypoxia-responsive elements (HREs) within the *RORA* and *KCTD16* promoters

Based on the previous data that PX478 attenuates the expressions of both RORA and KCTD16, we wonder if these two transcripts are also regulated by HIF1α since stabilization of HIF1α is a prerequisite for cellular adaptation to hypoxia and has been extensively documented in the literature (Semenza 2001). HIF-1α, a member of the basic helix-loop-helix Per-Arnt-Sim (bHLH-PAS) family of transcription factors (Wang et al. 1995), regulates the expression of a broad spectrum of genes collectively referred to as hypoxia-responsive genes (HRGs).

Given that *RORA* and *KCTD16* are upregulated under CoCl_2_ induced hypoxia and that HIF1α depletion significantly reduces their expression (Fig 4), we hypothesized that their promoters might contain HRE sequences bound by HIF1α. To investigate this, we reanalysed publicly available HIF1α ChIP-Seq data from hypoxic U2OS cells (GSM2257670) using ChIP-Atlas (https://chip-atlas.org/) and visualized the results in IGV (https://igv.org/). This analysis revealed clear HIF1α enrichment at the promoter/enhancer regions of both *RORA* and *KCTD16* (Fig 5 A and G). Further analysis of chromatin activation marks revealed that H3K4me3 (GSM4194654) and H3K27ac (GSM5488550) peaks overlapped with the HIF1α-enriched regions (Fig 5 A and G). Consistently, ATAC-Seq (GSM5899259) and RNA Pol II ChIP-Seq (GSM6439581) data demonstrated chromatin accessibility and transcriptional activity at these loci (Fig 5A and G). Collectively, these *in silico* results indicate that HIF1α binds to transcriptionally active regions within the *RORA* and *KCTD16* promoters under hypoxic conditions.

**FIGURE 5.**
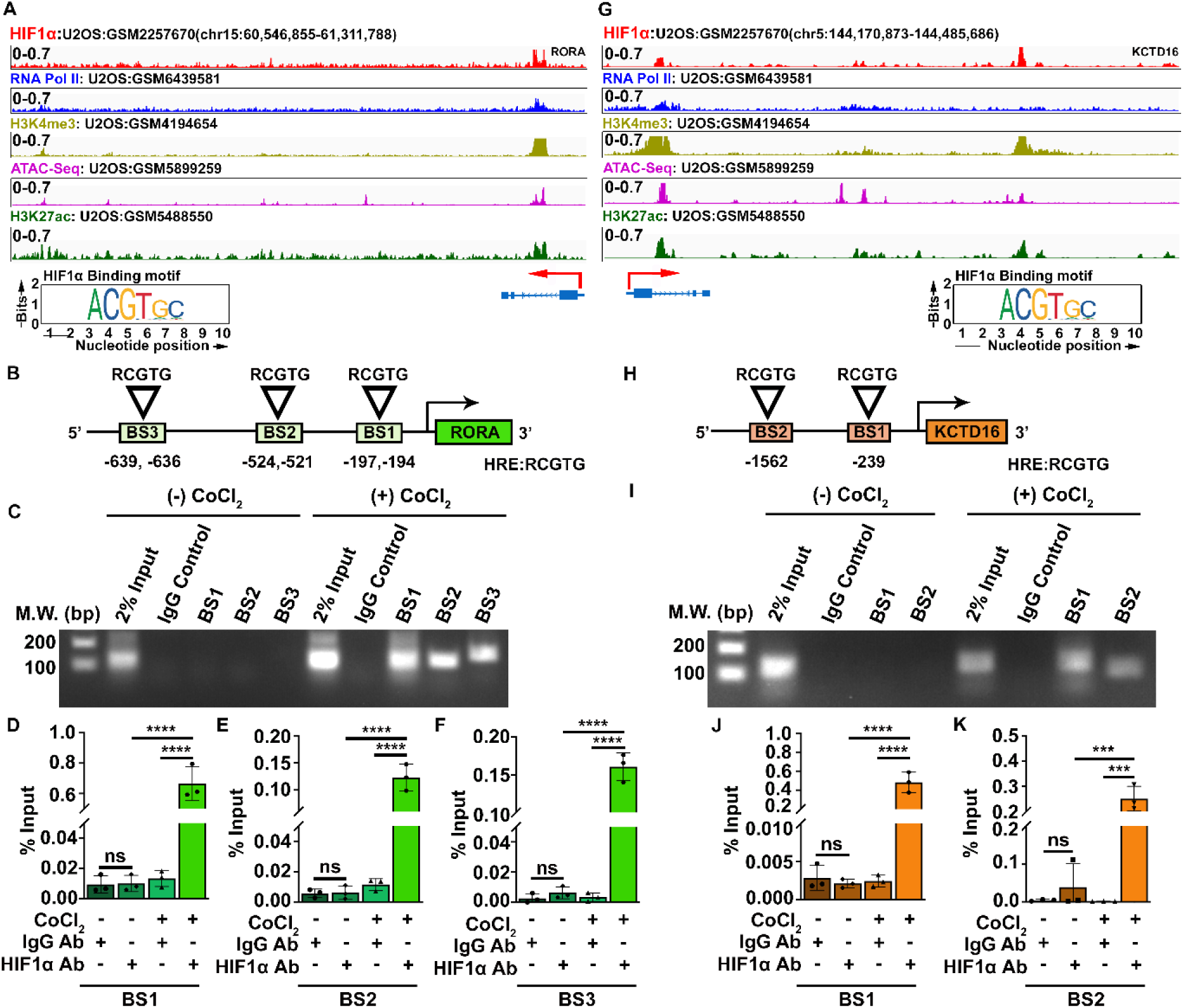
HIF1α binds in the promoter of *RORA* and *KCTD16*. (*A* and *G*) ChIP-Seq data from ChIP-Atlas visualized in IGV, showing HIF1α enrichment at chromatin-accessible regions of the RORA and KCTD16 promoters, respectively. (*B* and *H*) Schematic representation of predicted HIF1α binding sites (HREs) on the *RORA* and *KCTD16* promoters. (*C* and *I*) ChIP assays confirming direct HIF1α binding at *RORA* (BS1, BS2, BS3) and *KCTD16* (BS1, BS2) promoters; molecular markers indicated on the left. (*D*–*F*) ChIP-qPCR quantification of HIF1α occupancy at each binding site in the RORA promoter. (*J* and *K*) ChIP-qPCR quantification of HIF1α binding at BS1 and BS2 of the *KCTD16* promoter. Normalization was performed using the % input method. Data represent mean ± SD of three biological replicates; statistical significance was calculated using one-way ANOVA. ns: non-significant, *P < 0.05, **P < 0.005, ***P < 0.0005, ****P < 0.0001.

To identify specific HIF1α binding motifs, we performed *in silico* HRE prediction using the JASPAR transcription factor binding database (http://www.jaspar.genereg.net), focusing on the 5 kb upstream of each transcription start site. This analysis identified three clusters of closely spaced or overlapping HRE motifs in the *RORA* promoter (Fig 5B) and two putative HREs within the *KCTD16* promoter/enhancer region (Fig 5H) .To experimentally validate the *in silico* predictions, ChIP assays were performed using a HIF-1α-specific antibody in CoCl_2_ treated and untreated U2OS cells, followed by PCR amplification with primers targeting the predicted HREs. The results confirmed HIF1α binding to all three predicted clusters in the *RORA* promoter (Fig 5C) and to both predicted sites in the *KCTD16* promoter (Fig 5I) exclusively under CoCl_2_ induced hypoxia. No enrichment was observed with control IgG, and HIF1α binding was absent under normoxia, consistent with oxygen-dependent HIF1α degradation (Jaśkiewicz et al. 2022). Quantitative ChIP-qPCR using the % input method revealed differential enrichment across sites. In *RORA*, BS1 showed the highest HIF1α occupancy, followed by BS3 and BS2 (Fig 5D-F) whereas in *KCTD16*, BS1 was more enriched than BS2 (Fig 5J-K).

Together, these findings validate the *in-silico* predictions and demonstrate that HIF1α directly binds HREs within the promoter/enhancer regions of *RORA* and *KCTD16* under CoCl_2_ induced hypoxia. Combined with earlier observations that HIF1α depletion reduces expression of both genes (Fig 4), this establishes *RORA* and *KCTD16* as direct transcriptional targets of HIF1α and key components of the hypoxia-responsive transcriptome.

### *RORA* and *KCTD16* reduce osteosarcoma cellular aggressiveness by suppressing the levels of the cellular proliferation factor c-Myc under CoCl_2_ induced hypoxia

Our earlier results showed that CoCl_2_ induced hypoxia suppresses the aggressive phenotype of osteosarcoma cells. To uncover the underlying mechanism, we focused on c-Myc, a well-established proto-oncogene that is central to maintaining oncogenic properties in cancer cells (Melnik et al. 2019). Notably, previous studies have highlighted the importance of the c-Myc-HIFα signaling axis in regulating cellular adaptation to hypoxic stress (Huang 2008). Based on these reports, we examined c-Myc expression under CoCl₂-induced hypoxia. Immunoblot analysis revealed a pronounced reduction in c-Myc protein levels in hypoxic U2OS cells (Fig. 6 A and B), suggesting that hypoxia-mediated downregulation of c-Myc contributes to the diminished oncogenic potential observed under these conditions.

**FIGURE 6.**
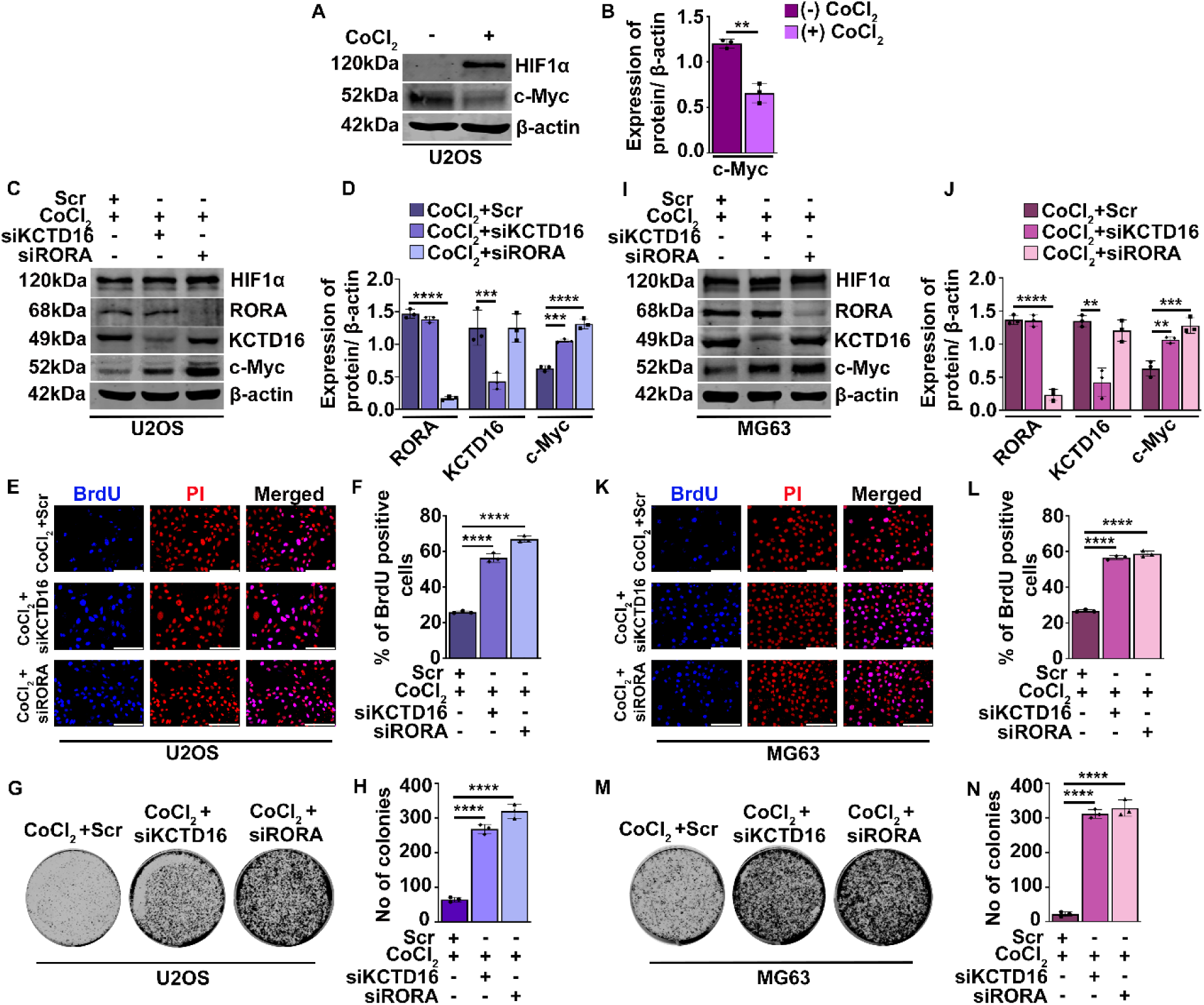
*RORA* and *KCTD16* augmented osteosarcoma cellular aggressiveness by elevating c-Myc expression. (*A*) Western blot analysis of c-Myc protein levels in U2OS cells under normoxic and CoCl₂-induced hypoxic conditions. (*B*) Densitometric quantification of c-Myc expression from panel A using ImageJ, showing reduced c-Myc levels in hypoxic U2OS cells. β-Actin served as the loading control. Statistical significance was determined using two-tailed Student’s *t*-test. (*C* and *E*) Western blots showing siRNA-mediated depletion of RORA (*C*) and KCTD16 (*I*) in U2OS and MG63 cells under hypoxic conditions. HIF1α blot confirms hypoxia induction. c-Myc levels were elevated upon depletion of either gene. (*D* and *J*) ImageJ-based densitometric quantification of blots from panels *C* and *I*, normalized to β-actin. Statistical analysis was performed using one-way ANOVA. (*E* and *K*) Representative microscopic images of BrdU incorporation assays in RORA and KCTD16 depleted hypoxic U2OS and MG63 cells, respectively. Scale bar = 50 μm. (*F* and *L*) Quantification of BrdU-positive cells showing increased proliferation upon RORA or KCTD16 depletion under hypoxic conditions (n ≥ 20 cells per condition). (*G* and *M*) Representative images of colony formation assays in RORA-and KCTD16-depleted hypoxic osteosarcoma cells. (*H* and *N*) Quantitative analysis showing enhanced clonogenic potential following RORA or KCTD16 depletion in hypoxic cells. Statistical significance was calculated using one-way ANOVA. All the data is represented as ± SD from three biological replicates. ns: non-significant, *P < 0.05, **P < 0.005, ***P < 0.0005, ****P < 0.0001.

To assess whether the hypoxia-responsive, cCE-targeted transcripts *RORA* and *KCTD16* influence c-Myc regulation, we performed siRNA-mediated knockdown of each gene in osteosarcoma cells under CoCl_2_ induced hypoxic conditions. Western blot analysis demonstrated that depletion of either RORA or KCTD16 led to a marked increase in c-Myc expression in both U2OS and MG63 cells, regardless of hypoxia induction (Fig 6 C and I). Efficient knockdown of RORA and KCTD16 was confirmed by immunoblotting, and elevated HIF1α levels verified successful hypoxia induction in CoCl_2_ treated samples (Fig 6 C and I). Densitometric analysis further revealed that while knockdown of both genes enhanced c-Myc levels, depletion of RORA caused a more pronounced increase in c-Myc expression than KCTD16 knockdown in both cell lines (Fig 6 D and J).

To corroborate the loss-of-function studies, EGFP-RORA and HA-KCTD16 were ectopically overexpressed in U2OS and MG63 cells. Immunoblotting with anti-EGFP and anti-HA antibodies confirmed robust expression of both constructs (Fig 7 *A*-*B* and *I*-*J*). Notably, overexpression of either RORA or KCTD16 led to a marked reduction in c-Myc levels, indicating an inverse correlation between RORA/KCTD16 abundance and c-Myc expression. Earlier studies showed that elevated c-Myc expression drives cellular aggressiveness in several cancers (Wang et al. 2008; Melnik et al. 2019; Dhanasekaran et al. 2022; Qiu et al. 2022). To investigate whether the elevated c-Myc levels resulting from RORA or KCTD16 depletion under CoCl_2_ induced hypoxia affect osteosarcoma aggressiveness, we performed two functional assays. First, the BrdU incorporation assay was used to quantify actively proliferating cells. Second, the colony formation assay assessed the ability of single cells to proliferate and form colonies of ≥50 cells. Both assays were carried out in CoCl_2_ induced hypoxic U2OS and MG63 cells following siRNA-mediated knockdown of RORA or KCTD16.

**FIGURE 7.**
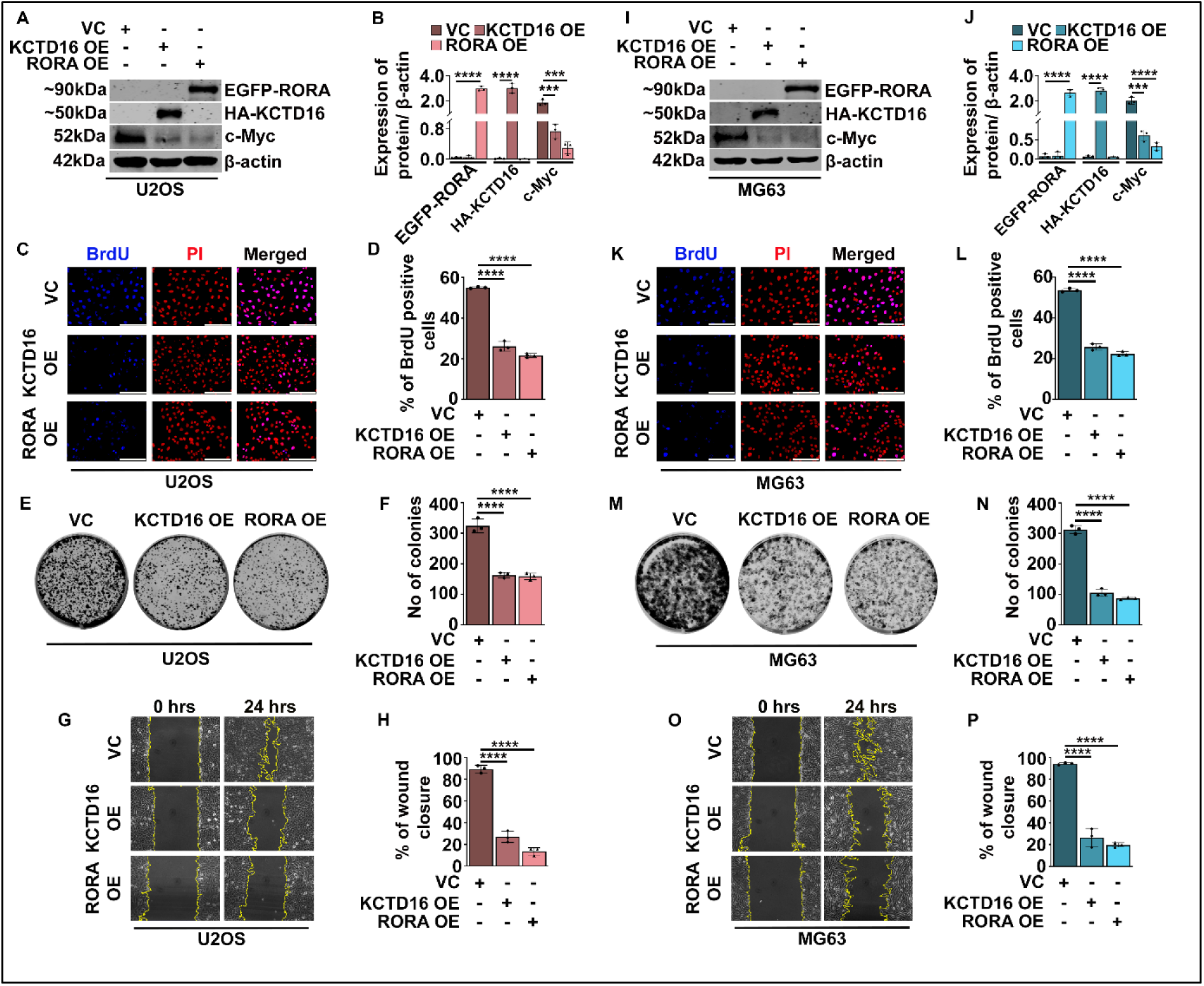
Precocious overexpression of either RORA or KCTD16 suppresses osteosarcoma cellular aggressiveness by downregulating c-Myc expression. (*A* and *I*) Western blots showing RORA, KCTD16, and c-Myc levels in U2OS and MG63 cells, respectively. (*B* and *J*) Densitometric quantification of the blots using ImageJ demonstrated that overexpression of RORA or KCTD16 led to reduced c-Myc expression in both cell lines. β-Actin was used as a loading control for normalization. (*C* and *K*) Representative microscopic images of BrdU incorporation assays in RORA and KCTD16 overexpressing U2OS and MG63 cells, respectively. Scale bar = 50 μm. (*D* and *L*) Quantification of BrdU-positive cells showing significantly decreased proliferative capacity in RORA and KCTD16-overexpressing cells (n ≥ 20 cells per condition). (*E* and *M*) Representative images of colony formation assays in RORA and KCTD16 overexpressing osteosarcoma cells. (*F* and *N*) Quantification of colonies demonstrating a marked reduction in clonogenic potential upon RORA or KCTD16 overexpression. (*G* and *O*) Representative images of migration assays in RORA and KCTD16-overexpressing cells. Scale bar = 50 μm. (*H* and *P*) Quantification of migrated cells showing significantly impaired migratory capacity in RORA and KCTD16 overexpressing osteosarcoma cells. Statistical analysis was performed using one-way ANOVA, and all data are presented as mean ± SD from three independent biological replicates. Significance is indicated as follows: ns, not significant; *P < 0.05; **P < 0.005; ***P < 0.0005; ****P < 0.0001.

Depletion of RORA or KCTD16 in CoCl_2_ treated U2OS and MG63 cells markedly increased both proliferative capacity and clonogenic potential compared with scrambled-transfected hypoxic controls (Fig 6 *E*-*F* and *K*-*N*). In contrast, ectopic overexpression of either transcript significantly suppressed proliferation, colony formation, and migratory ability (Fig 7 *C*-*H* and *K*-*P*). These findings align with previous reports, such as miR-652-mediated repression of RORA promoting proliferation and metastasis in endometrial cancer (Sun et al. 2018). Importantly, our study provides the first evidence that hypoxia-responsive, cCE targeted RORA acts as a suppressor of osteosarcoma aggressiveness under CoCl_2_ induced hypoxia, and it also identifies KCTD16 as a novel HRR family member with tumor-suppressive function in CoCl_2_ induced hypoxic osteosarcoma cells. Together, these results demonstrate that the upregulation of RORA and KCTD16 duing CoCl_2_ induced hypoxia is essential for limiting osteosarcoma cell aggressiveness, highlighting their key role as negative regulators of hypoxia-driven tumor progression.

Some studies have reported that other KCTD family members, such as KCTD2, can function as adaptor proteins that directly interact with c-Myc and promote its destabilization by recruiting the Cullin3 E3 ubiquitin ligase complex through their BTB domain(Kim et al. 2017). Notably, KCTD16 also contains a BTB domain; however, there is currently no evidence supporting its interaction with c-Myc. To investigate this possibility, we performed *in silico* protein-protein docking analysis using ClusPro. The docking results suggested a potential interaction between KCTD16 and c-Myc (Supplemental Fig. 3A-B). To validate this finding, we conducted an HA-KCTD16 pull-down assay. Immunoprecipitation using anti-HA revealed the presence of c-Myc as a co-immunoprecipitated protein, thereby confirming a physical interaction between HA-KCTD16 and c-Myc (Supplemental Fig. 3C). These findings suggest the possibility that KCTD16, via its BTB domain, interacts with c-Myc and, potentially in association with an as-yet unidentified factor, promotes its degradation.

## DISCUSSION

Cells encountering unfavourable environmental conditions initiate specialized stress-response mechanisms to maintain cellular homeostasis. Hypoxia represents one such hostile microenvironment, defined by low oxygen levels (≤1%) and commonly observed in solid tumours (Abou Khouzam et al. 2023; Chen et al. 2023). As the master regulator of the hypoxic response, HIF1α reshapes the cellular transcriptomic landscape by modulating the expression of numerous hypoxia-responsive genes (HRGs), thereby initiating hypoxia-adaptive signalling pathways (Yfantis et al. 2023). Increasing evidence indicates that intratumoral hypoxia promotes cancer progression through the activation of diverse intracellular signalling cascades (Kunachowicz et al. 2025; Rahman et al. 2026). Conversely, several studies have reported that hypoxia can also attenuate cellular aggressiveness (Gardner et al. 2001; Kumar and Vaidya 2016; Shi et al. 2023). A careful evaluation of the existing literature suggested that the cellular response to hypoxia is highly context dependent. These determinants include (i) the specific cell type, (ii) the method of hypoxia induction, such as physical oxygen deprivation versus chemical mimetics, (iii) the duration of hypoxic exposure, and (iv) the concentration of hypoxia-mimicking agents (Jaśkiewicz et al. 2022; Zhang et al. 2026). Supporting this view, Calvo-Anguiano *et al*. demonstrated that transcriptomic profiles generated under distinct hypoxic paradigms exhibit overlapping yet divergent sets of regulated genes, indicating that hypoxic responses are not uniform but are instead shaped by the nature of the hypoxic stimulus (Calvo-Anguiano et al. 2018). Furthermore, another study reported a dose-dependent effect of CoCl_2_ on breast cancer cells, wherein cellular proliferation increased progressively up to a concentration of 150 µM, followed by a reduction in proliferation at higher doses (Zhang et al. 2013).

Given that our work was confined to an *in vitro* chemically induced hypoxia model, we initially investigated the effects of CoCl_2_ induced hypoxia on osteosarcoma cells. Based on this approach, osteosarcoma cells were exposed to 200 µM CoCl_2_ for 24 hrs. Consistent with previous findings reported by Zhang *et al*. (2013), treatment with 200 µM CoCl_2_ significantly reduced osteosarcoma cell aggressiveness, as evidenced by diminished proliferative capacity, colony formation, and migratory ability. Furthermore, cell-cycle progression was disrupted, with a reduced proportion of cells in the S phase and an increased accumulation in the G₂/M phase relative to control cells. Importantly, treatment with 200 µM CoCl_2_ did not adversely affect overall cell viability, in agreement with our earlier observations (Islam and Mukherjee 2025).

Beyond HIF-mediated transcriptional regulation, recent studies emphasize the importance of post-transcriptional mechanisms in cellular adaptation and survival under hypoxic conditions (Oltean and Bates 2014; Gallego-Paez et al. 2017; Khabar 2017). Among these regulatory processes, cytoplasmic capping enzyme (cCE) mediated recapping of target transcripts (including both coding and noncoding RNAs) has garnered considerable attention in recent years (Mukherjee et al. 2012; Kiss et al. 2016; Mukherjee et al. 2023; Islam and Mukherjee 2025). The present study demonstrates that CoCl_2_ induced hypoxia attenuates the aggressiveness of osteosarcoma cells. Given the absence of detailed molecular insights in the existing literature regarding the suppression of osteosarcoma cell aggressiveness by 200 µM CoCl_2_ induced hypoxia, we sought to elucidate the underlying molecular mechanisms. Transcriptomic profiling revealed a significant upregulation of *RORA* and *KCTD16* in osteosarcoma cells subjected to CoCl_2_ induced hypoxic conditions. Studies showed that the HIF1α-HIF1β heterodimer binds to hypoxia-responsive elements (HREs), typically the consensus motif 5′-RCGTG-3′, located in the promoter or enhancer regions of target genes and, with the assistance of transcriptional co-activators, activates their transcription (Semenza et al. 1997). Similarly, our analyses showed that the hypoxia-driven induction of *RORA* and *KCTD16* is mediated by HIF1α. Functionally, elevated levels of these hypoxia-responsive transcripts suppressed the expression of the proliferative regulator c-Myc, resulting in reduced cellular aggressiveness, as evidenced by decreased proliferation, colony formation, migration, and invasion capacities (Fig. 8). Consistent with these observations, hypoxic stress also induced cell-cycle arrest. Notably, this study further demonstrates that *RORA* and *KCTD16* are post-transcriptionally stabilized by cCE, as inhibition of cCE activity using the K294A mutant led to a marked reduction in their expression. In agreement with our previous study (Islam and Mukherjee 2025), cCE maintains the stability of these transcripts during CoCl_2_ induced hypoxia too. Functional perturbation experiments revealed that silencing either *RORA* or *KCTD16* enhanced osteosarcoma cell aggressiveness, whereas their overexpression produced the opposite effect.

**FIGURE 8.**
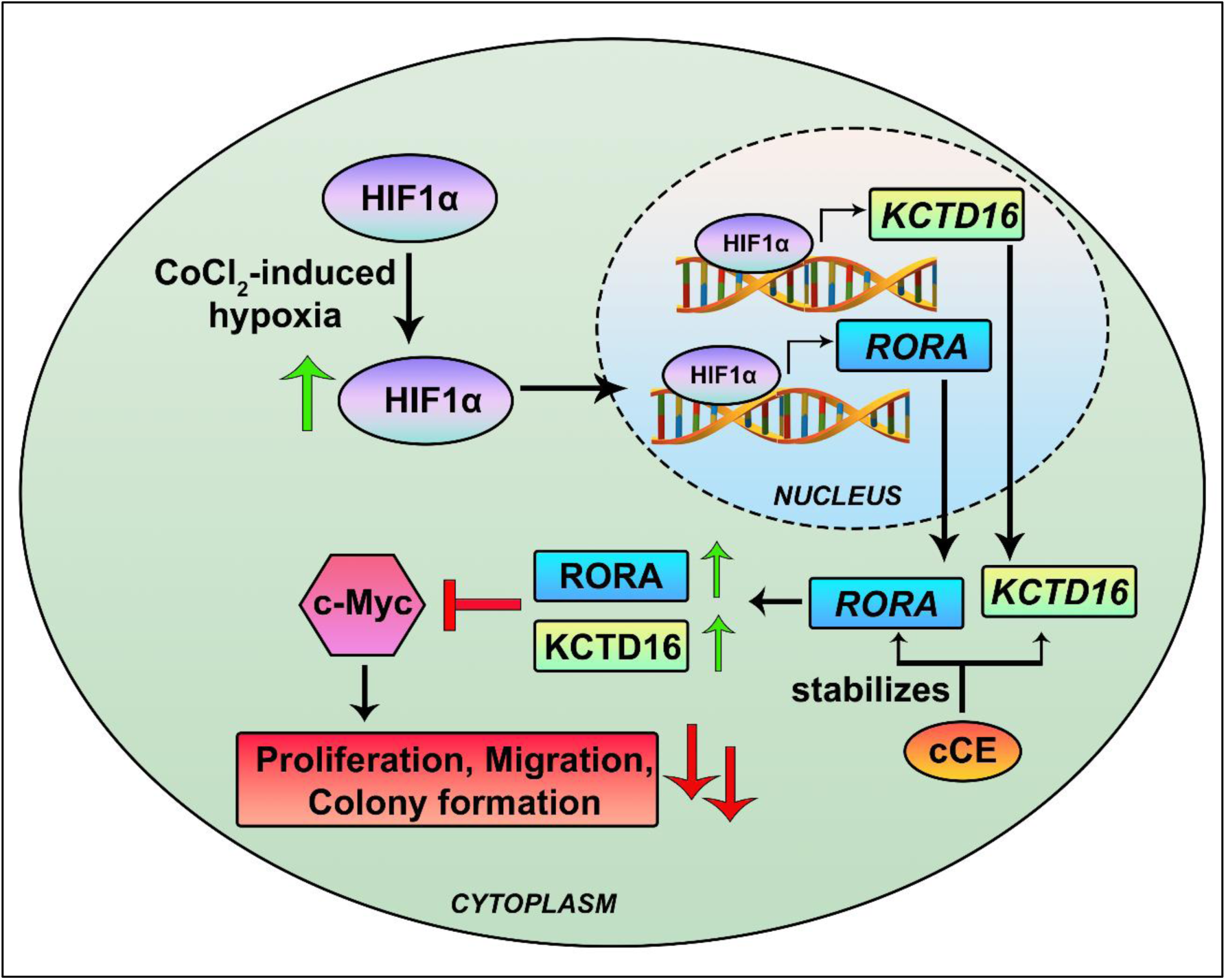
Schematic illustration depicting the impact of CoCl_2_ induced hypoxia on osteosarcoma cells. CoCl_2_ treatment stabilizes HIF1α, leading to the transcriptional upregulation of the hypoxia-responsive RNAs *RORA* and *KCTD16*. These transcripts are subsequently stabilized at the post-transcriptional level by cCE. cCE-mediated protection of these hypoxia-responsive RNAs HRRs attenuates hypoxia-induced osteosarcoma aggressiveness through suppression of the key proliferative factor c-Myc.

Transcriptomic profiling of hypoxic osteosarcoma cells identified a large number of differentially expressed genes. Focusing specifically on hypoxia-responsive pathways, we selected *RORA* and additionally identified *KCTD16* as a prominently upregulated transcript under hypoxic conditions. Notably, a comprehensive literature survey revealed no established association between *KCTD16* and cancer. However, pan-cancer analysis using The Cancer Genome Atlas (TCGA) demonstrated that *KCTD16* expression is downregulated across several solid tumours. Similarly, TCGA datasets indicated that *RORA* expression is reduced in the majority of solid tumours.

Consistent with these observations, previous studies have reported that RORA suppresses cancer cell aggressiveness in glioma, gastric cancer, and endometrial cancer (Sun et al. 2018; Jiang et al. 2020; Ma et al. 2021). However, the role of RORA in osteosarcoma is yet to be known. Recent evidence suggests that RORA attenuates c-Myc expression by transcriptionally activating the ubiquitin ligase NEDD4, which promotes c-Myc ubiquitination and subsequent proteasomal degradation (Wang et al. 2022). However, our RNA-seq analysis did not reveal any alteration in NEDD4 expression, indicating that RORA may regulate c-Myc through alternative direct or indirect mechanisms in hypoxic osteosarcoma cells. Elucidation of these mechanisms warrants further investigation.

In contrast, the role of KCTD16 in cancer biology remains unexplored. Nevertheless, other members of the KCTD family have been implicated in tumour suppression; for example, KCTD2 inhibits glioma progression by facilitating c-Myc degradation via interaction with the Cullin3 E3 ubiquitin ligase through its BTB domain, while KCTD10 suppresses lung and colorectal cancer development (Kim et al. 2017; Yin et al. 2025a; Yin et al. 2025b). Although KCTD16 shares structural similarity with KCTD2, including the presence of a BTB domain, emerging studies indicate that KCTD16 is unable to form a stable complex with Cullin3 (Pinkas et al. 2017) but our data showed the potential interaction between KCTD16 and c-Myc. This raises an important mechanistic question regarding how KCTD16 suppresses c-Myc expression in the absence of Cullin3 interaction. Addressing this question will be a critical focus of future studies aimed at defining KCTD16-dependent regulatory pathways in hypoxic osteosarcoma. Functionally, our data demonstrate that silencing either *RORA* or *KCTD16* under CoCl_2_ induced hypoxic conditions significantly enhances cellular aggressiveness, whereas ectopic overexpression of either gene markedly reverses this phenotype. These findings confirm the tumour-suppressive role of RORA, in agreement with prior studies, and establish a direct link between KCTD16 and hypoxia.

As this study primarily relies on CoCl_2_ induced hypoxia, which mimics hypoxic signalling through HIF1α stabilization but does not fully recapitulate physiological hypoxic conditions, future investigations employing physical hypoxia models are warranted. Additionally, our findings suggest that HRRs can serve as substrates for cytoplasmic capping enzyme (cCE), underscoring the need for a systematic exploration of the hypoxic transcriptome to identify additional HRRs regulated by cCE-mediated recapping. Collectively, this study identifies *RORA* and *KCTD16* as integral components of the hypoxic regulatory network and reveals a previously underappreciated role for cCE in modulating tumour-suppressive pathways under hypoxic stress.

## MATERIALS AND METHODS

### 3.1 Cell culture and treatments

Human osteosarcoma cell lines U2OS and MG63 were procured from the American Type Culture Collection (ATCC) and the National Centre for Cell Science (NCCS), Pune, respectively. All the cell lines were cultivated in 1X Dulbecco’s Modified Eagle’s Media (DMEM) (Gibco; cat# 12100046;) fortified with 10 % fetal bovine serum (FBS) (Himedia; cat# RM1112) and a cocktail of penicillin-streptomycin (Gibco; cat# 15140-122) to a final concentration of 1 %, in a humidified CO_2_ incubator (Eppendorf) where the concentration of CO_2_ was set at 5 % and temperature was maintained at 37 °C. Cells were exposed to different treatments for experimentation upon reaching a confluency of 60-70 %. A mycoplasma contamination check was routinely done using a PCR detection kit (Sigma-Aldrich; cat# MP0035) to ensure contamination-free cell culture.

An *in vitro* chemically induced hypoxic model was established as discussed in using CoCl_2_ (Sigma Aldrich; cat no# C8661) (Islam and Mukherjee 2025). Briefly, 70-80 % confluent osteosarcoma cells (U2OS or MG63) were treated with CoCl_2_ solution prepared in DPBS (stock concentration 25 mM) to a final concentration of 200 μm for 24 hrs. Following incubation, cells were harvested for RNA and protein isolation.

An *in vitro* HIF-1α inhibition assay was conducted by treating osteosarcoma cells with the potent HIF-1α inhibitor PX478 (MedChemExpress; cat# HY-10231) at a final concentration of 40 μM, either in the presence or absence of CoCl_2_ for 24 hrs as done previously (Islam and Mukherjee 2025).

### 3.2 Cloning of human full-length coding region (CDS) *RORA* and *KCTD16*

U2OS whole cell RNA was reverse transcribed using the SuperScript III First-Strand cDNA Synthesis Kit (Invitrogen; cat# 18080051) following manufacturers instruction. The full-length wild-type coding sequences of *RORA* (NM_134261.3) and *KCTD16* (NM_001370487.1) were PCR-amplified from U2OS whole-cell cDNA using Q5 High-Fidelity DNA Polymerase (NEB; cat# M0491L) according to the manufacturer’s protocol. The primers used for amplification are listed in Table 1. The amplified *RORA* cDNA was inserted into the pcDNA4/TO-EGFP plasmid at the BamHI and NotI sites using the In-Fusion cloning strategy (Takara Bio; cat# 638909). For *KCTD16* cloning, during *KCTD16* amplification, a forward primer encoding an N-terminal HA tag (Table 1) was employed to generate an HA-tagged KCTD16 amplicon. The resulting HA-KCTD16 cDNA was cloned into the pcDNA4/TO vector at the PstI and XbaI sites using the In-Fusion cloning system (Takara Bio). Briefly, the pcDNA4/TO-EGFP vector used for RORA cloning was linearized with BamHI and NotI, while the pcDNA4/TO vector used for KCTD16 cloning was linearized with PstI and XbaI. The digested vectors were gel-purified using the NucleoSpin Gel and PCR Clean-up Kit (Macherey-Nagel; cat# 740609.250). The corresponding inserts were purified using the same kit. Purified vector and insert were mixed at a 1:2 molar ratio and subjected to the In-Fusion cloning reaction by incubation at 50 °C for 15 min. The reaction products were directly transformed into chemically competent DH5α cells, and positive clones were selected on ampicillin-containing Luria-Bertani agar plates. Initial screening was performed by restriction digestion using BamHI/NotI (for RORA) or PstI/XbaI (for KCTD16) to confirm insert drop-out. Positive clones were further validated by digestion with additional restriction enzyme combinations, followed by Sanger sequencing to confirm insert integrity and sequence accuracy.

**Table 1:**
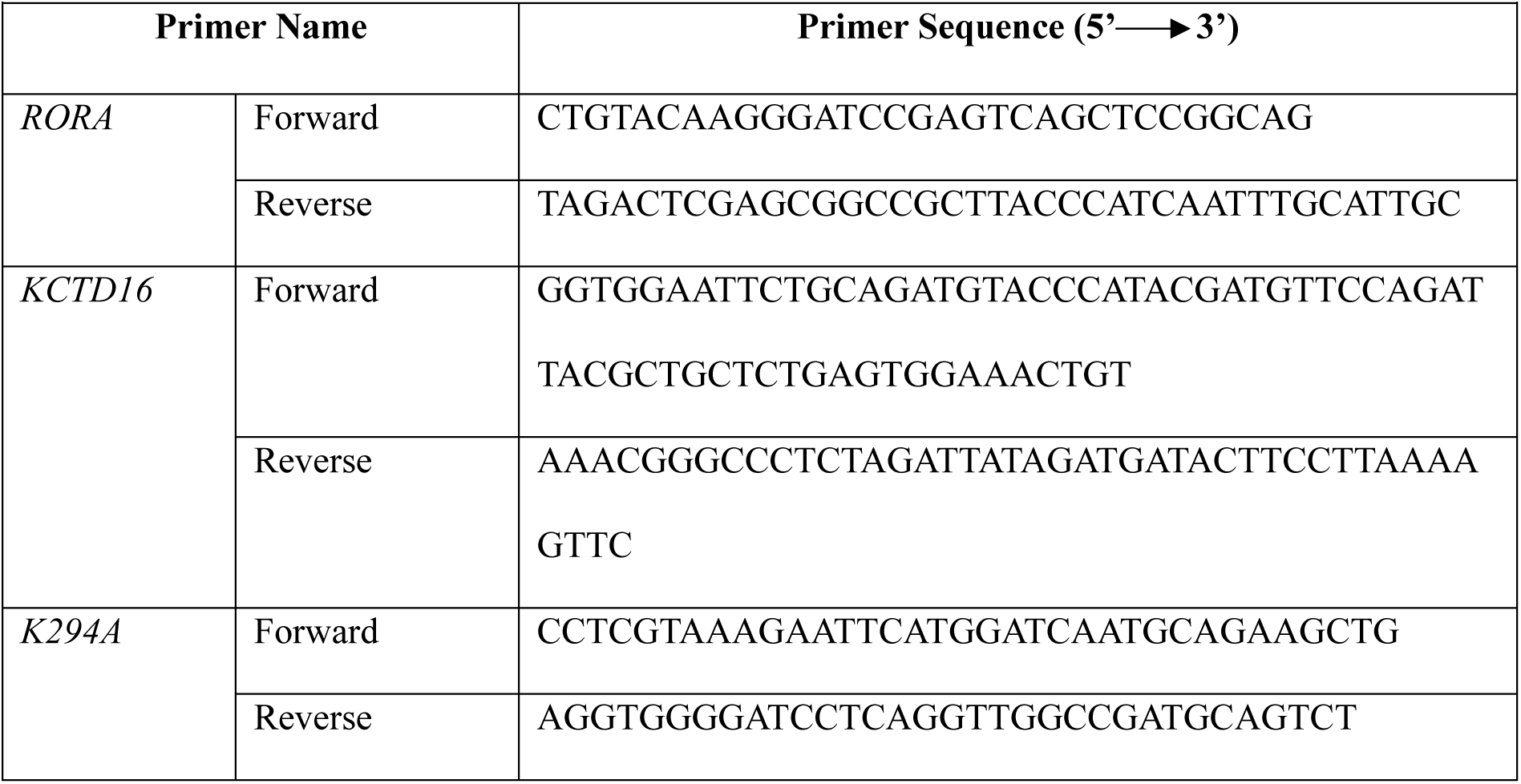
List of cloning primers used in the study.

### 3.3 Generation of a stable cell line expressing cytoplasmically restricted catalytically inactive Capping Enzyme (K294A)

The cloning of pLVX-TetOne-Puro myc-mCE(K294A) and the generation of the stable cell line expressing myc-K294A have been described previously (Islam and Mukherjee 2025). Briefly, the mouse cytoplasm-restricted, catalytically inactive capping enzyme was PCR-amplified from pcDNA4/TO-bio-myc-NES-mCE(K294A)ΔNLS using Q5 High-Fidelity DNA Polymerase (NEB; cat# M0491L) and subcloned into the pLVX-TetOne-Puro vector (Takara Bio; cat# 631847) through homologous recombination using the In-Fusion cloning system (Takara Bio; cat# 638909). For this purpose, the vector was linearized with BamHI and EcoRI restriction enzymes. The primer sequences are listed in Table 1.

For an inducible stable cell line expressing myc-mCE(K294A), U2OS cells at 60–70% confluency were transfected with pLVX-TetOne-Puro-myc-K294A using JetPRIME transfection reagent (Polyplus; cat# 101000046) according to the manufacturer’s instructions. After 48 h, cells were diluted and selected with puromycin (50 μg/ml; Sigma-Aldrich; cat# 9620) until single clones were isolated. Selected clones were induced with doxycycline (1 μg/ml; Sigma-Aldrich; cat# D5207), and clones showing the highest myc-mCE(K294A) expression were used for subsequent experiments.

### 3.4 Biochemical fractionation into cytoplasmic and nuclear fractions

To prepare cytoplasmic lysates from osteosarcoma cells (U2OS and MG63), cells were first harvested in 1× DPBS and harvested by centrifugation at 1,000*g* for 5 min at 4 °C. The supernatant was discarded, and the cell pellet was resuspended in 1X cytoplasmic lysis buffer (20 mM Tris-Cl, pH 7.5; 10 mM NaCl; 10 mM MgCl₂; 10 mM KCl; 0.2% NP-40) supplemented with 1X protease inhibitor cocktail (Roche; cat# 12352204), 1 mM PMSF (Thermo Scientific; cat# 36978), and RNaseOUT (Invitrogen; cat# 10777019). The cell suspension was incubated on ice for 10 min with gentle tapping. Following incubation, samples were centrifuged at 1,000*g* for 10 min at 4 °C, and the supernatant was collected as the cytoplasmic lysate.

The nuclear pellet obtained during cytoplasmic fractionation was washed with 1X DPBS to eliminate residual cytoplasmic contamination. The washed nuclear pellet was then resuspended in 1X nuclear lysis buffer (150 mM NaCl, 1% Triton X-100, 0.5% sodium deoxycholate, 1% SDS, and 25 mM Tris-Cl, pH 7.5) supplemented with 1X protease inhibitor cocktail (Roche; cat# 12352204), 1 mM PMSF (Thermo Scientific; cat# 36978), and RNaseOUT (Invitrogen; cat# 10777019). The suspension was incubated on ice for 40 min with vigorous agitation. Nuclear lysates were then collected by centrifugation at 10,000*g* for 10 min at 4 °C.

### 3.5 RNA isolation and poly(A) RNA preparation

Total RNA was extracted by directly adding TRIzol reagent (Invitrogen; cat# 155960260) to the cell pellet, whereas cytoplasmic RNA was isolated by adding TRIzol reagent to the cytoplasmic fraction obtained as described in the previous section, following the manufacturer’s instructions. Briefly, 1 mL of TRIzol containing the cellular suspension was mixed with 200 μL chloroform (Merck; cat# C2432) by vigorous vortexing and incubated at room temperature for 2 min. The samples were then centrifuged at 12,000*g* for 15 min at 4 °C to achieve phase separation. The aqueous phase was carefully transferred to a fresh tube, and RNA was precipitated by adding isopropanol (Sisco Research Laboratories Pvt. Ltd; cat# 38446) at a 1:1 ratio (aqueous phase: isopropanol), along with 2 μL GlycoBlue (Invitrogen; cat# AM9516). Following incubation at room temperature for 15-20 min, samples were centrifuged at 12,000*g* for 10 min at 4 °C. The resulting RNA pellet was washed with 75% ethanol (HiMedia; cat#MB228), air-dried, and resuspended in ultrapure distilled water. To eliminate genomic DNA contamination, RNA samples were treated with DNase I (Thermo Scientific; cat# EN0521) according to the manufacturer’s instructions. DNA-free RNA was subsequently purified using phenol: chloroform: isoamyl alcohol (Sisco Research Laboratories Pvt. Ltd; cat# 69031) following the manufacturer’s protocol.

Poly(A) RNA was enriched from total RNA using the Dynabeads mRNA DIRECT Kit (Invitrogen; cat# 61011) according to the manufacturer’s instructions. Briefly, 2 μg of DNA-free total RNA was incubated with oligo(dT)-conjugated Dynabeads at 25 °C for 30 min in a thermomixer. The RNA-bead complexes were sequentially washed with Wash Buffers A and B, and poly(A) RNA was eluted in 10 mM Tris-HCl (pH 7.5).

### 3.6 cDNA preparation and quantitative real time PCR (qPCR)

A total of 2 μg of whole-cell or cytoplasmic RNA was reverse transcribed using the SuperScript III First-Strand cDNA Synthesis Kit (Invitrogen; cat# 18080051) following the manufacturer’s instructions. Quantitative PCR was performed using SsoAdvanced Universal SYBR Green Supermix (Bio-Rad; cat# 1725271) on a CFX Connect Real-Time PCR System (Bio-Rad) under standard reaction conditions as discussed before with a 1:1 diluted cDNA as the template (Islam and Mukherjee 2025). *Rplp0* served as the internal control, and relative gene expression was calculated using the comparative Ct (ΔΔCt) method. At least three independent biological replicates were analysed unless stated otherwise. The list of the primers used in this study is tabulated in Table 2.

**Table 2:**
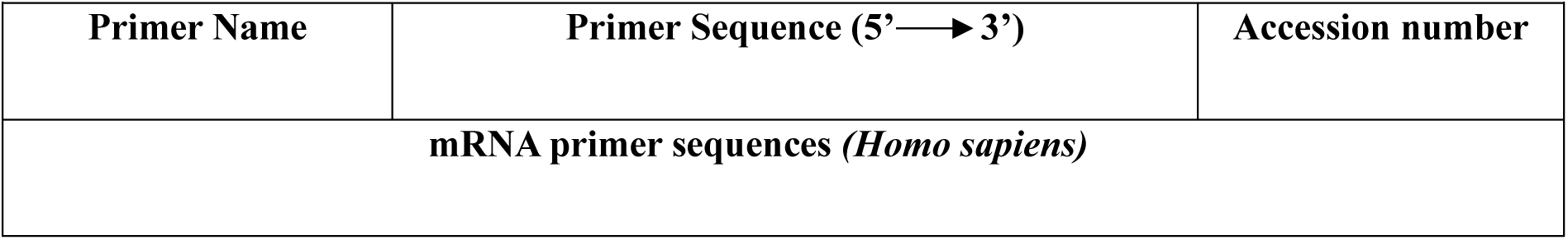
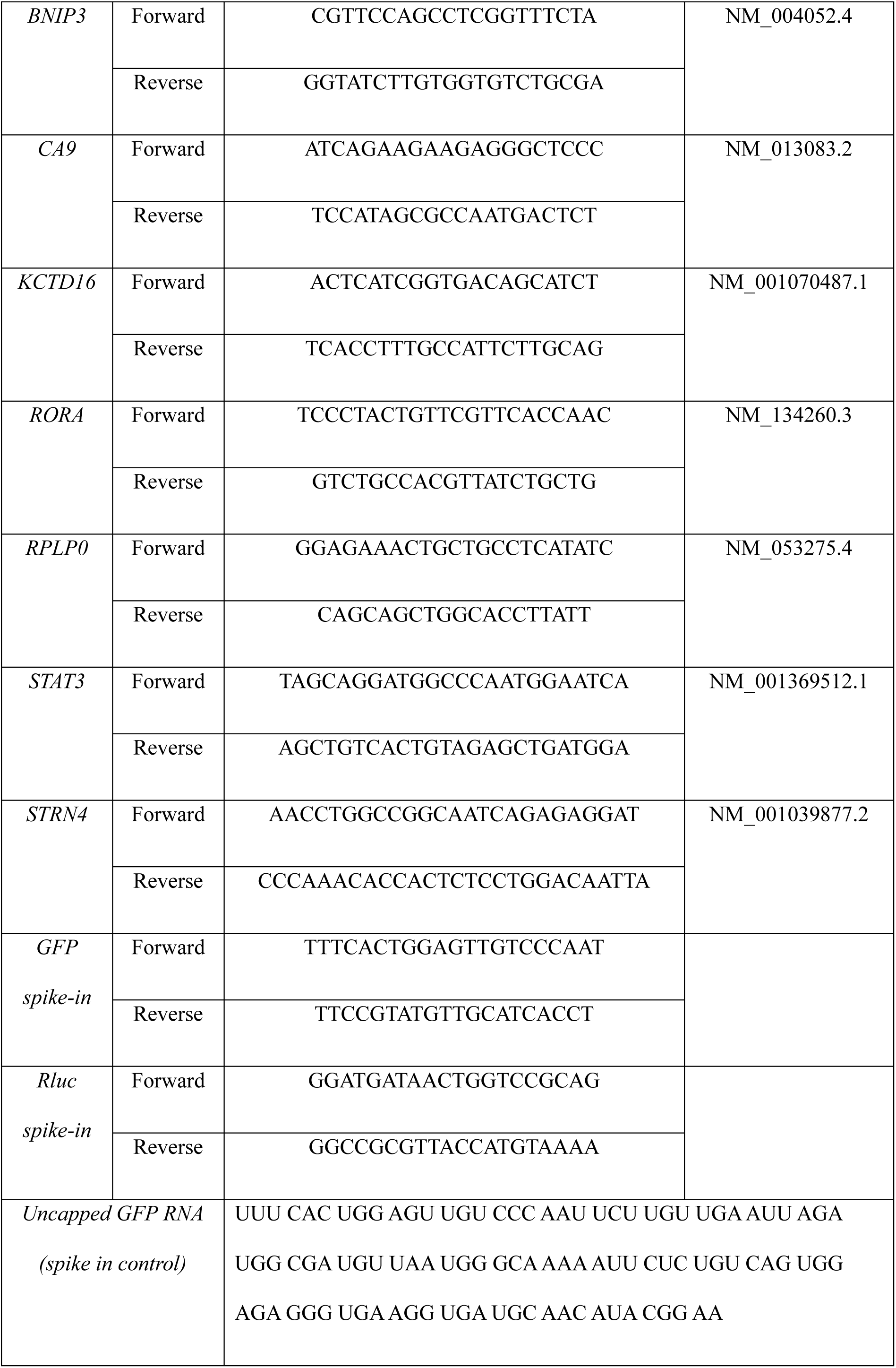
List of qPCR primers used in the study.

### 3.7 siRNA-mediated depletion of *Xrn1*, *KCTD16*, and *RORA* in osteosarcoma cells

For *Xrn1* knockdown, K294A stable cells were transfected with a predesigned siRNA targeting the coding region of *Xrn1*. Cells at 60–70% confluency were transfected with 10 nM siXRN1 (Ambion; Assay ID; s29016) using Lipofectamine RNAiMAX (Invitrogen; cat# 13778-075) according to the manufacturer’s instructions. The siRNA concentration was optimized to achieve maximal *Xrn1* knockdown. At 24 hrs post-transfection, the medium was replaced with fresh culture medium. At 48 hrs post-transfection, cells were treated with doxycycline (Dox; final concentration 1 μg/mL) to induce K294A expression or left untreated as controls. Cells were maintained for an additional 24 hrs and harvested 72 hrs post-transfection for RNA and protein isolation (Islam and Mukherjee 2025).

The expression of *KCTD16* and *RORA* was downregulated in CoCl_2_ treated hypoxic osteosarcoma cells. Briefly, either U2OS or MG63 cells were transfected with predesigned siRNAs specific for *KCTD16* (Ambion; Assay ID; s33231) or *RORA* (Ambion; Assay ID; s12105) at a 20 nM concentration using the method discussed above. At 48 hrs post-transfection, cells were treated with 200 μM CoCl₂ for 24 hrs to induce hypoxia. Following incubation, cells were harvested for RNA or protein extraction as required. Scrambled siRNA (scr) was used as a negative control in all siRNA-mediated knockdown experiments.

### 3.8 Ectopic over expression of *KCTD16*, and *RORA* in osteosarcoma cells

For ectopic overexpression of *KCTD16* and *RORA*, osteosarcoma cells (U2OS and MG63) were transfected with 1 μg of HA-KCTD16 or EGFP-RORA plasmid using JetPRIME transfection reagent (Polypus;, cat# 101000046) at a DNA: reagent ratio of 1:1, following the manufacturer’s instructions. The control cells were transfected with empty vectors. Briefly, 1 μg plasmid DNA was mixed with 200 μL transfection buffer, followed by the addition of 1 μL JetPRIME reagent, gently vortexed, briefly spun down, and incubated at room temperature for 10 min. The transfection mixture was then added dropwise to cells in fresh growth medium and gently mixed. At 24 hrs post-transfection, the medium was replaced with fresh culture medium. Cells were harvested 48 hrs post-transfection for RNA and protein isolation.

### 3.9 Western hybridization

Western blot was performed as discussed previously (Islam and Mukherjee 2025). Briefly, cells from different treatment conditions were harvested in 1X chilled DPBS and were centrifuged at 1000*g* for 5 min at 4 °C. Whole cell protein was isolated by lysed the cell pellet in 1X ice cold lysis buffer (50 mM Tris-Cl pH 8.0, 150 mM NaCl, and 1 % NP-40) fortified with1 mM PMSF and protease cocktail inhibitor (Roche; cat# 4693159001) and kept in ice for 30 min with intermittent vortexing. Once incubation was over, the whole cell protein was collected by centrifuging the lysed cells at 12000*g* for 10 min at 4 °C. The quantification of the protein was done by performing the BCA protein assay (Pierce; cat# A53226) following the instructions. Approximately 30-60 μg protein was resolved in 8-10% SDS-PAGE under reducing conditions and transferred onto Immobilon-FL PVDF (Millipore; cat# IPFL00010) membranes. All the membranes were blocked in 3 % skimmed milk in phosphate-buffered saline (PBS) for 30 min at room temperature, followed by incubation in the respective primary antibody dilution at 4 °C on the rocker overnight. The working dilutions of the antibodies used in the study are as follows 1:1000 rabbit anti-HIF1α (Cell Signaling Technology; cat# 36169); 1:1000 mouse anti-beta actin (Santa Cruz Biotechnology; cat# SC-517582), 1:2000 mouse anti-GAPDH (Novus Biologicals; cat# 2D4A7); 1:2000 rabbit anti-CA9 (Cloud-clone Crop; cat# PAD076Hu01); 1:2000 mouse anti-lamin (DHSB, cat# MANLAC1(4A7), 1:1000 rabbit anti-Xrn1 antibody, 1:1000 rabbit anti-KCTD16 (Proteintech; cat# 25437-1-AP), 1:1000 rabbit-anti-BNIP3 (Cloud-clone Crop; cat# PAJ545Ra01), 1:1000 rabbit anti-RORA (Cloud-clone Crop; cat# PAD947Hu01), 1:1000 mouse anti-c-myc (Santa Cruz Biotechnology; cat# SC-40). Following primary antibody incubation, membranes were washed thrice with PBS-T and incubated with respective secondary antibodies like donkey anti-MouseDyLight 800 (Cell Signalling Technology; cat no#5257) and donkey anti-Rabbit 680 (Invitrogen; cat# A10043) at 1: 10000 dilutions in PBS-T for 1 hr at room temperature. Following secondary antibody incubation membrane were again washed three times, and blots were developed using OdysseyCLx Imaging System (LiCor Inc.), and band intensities were quantified using ImageJ software (NIH, USA).

### 3.9 Cap Analysis of Gene Expression (CAGE)

Cap analysis of gene expression (CAGE) is a widely used technique for mapping transcription start sites (TSSs). Subsequent large-scale applications of CAGE in mammalian transcriptomes revealed that a subset of CAGE tags did not align with canonical TSSs but instead mapped within gene bodies(2009). Later studies suggested that transcripts harbouring such internal CAGE tags are likely substrates of the cytoplasmic capping machinery. To identify internal CAGE tags in selected mRNA transcripts, a computational pipeline was used to aggregate CAGE peaks relative to annotated genes and transcripts. This workflow quantifies local CAGE peak density across genomic loci by binning peaks into user-defined, non-overlapping windows of fixed size (e.g., 10 kb). For each gene or transcript, all mapped CAGE peaks were extracted, and their start positions were assigned to the corresponding genomic intervals. *Window* = (*peakstartw*)× *W*;, where W is the window size in base pairs. The frequency of peaks per window is then computed, producing a quantitative profile of transcription initiation activity across the region. These distributions were visualised using bar plots showing the number of CAGE peaks per genomic bin. This approach facilitates the identification of transcriptional hotspots, putative alternative promoters, and clusters of uncapped mRNAs, which are increasingly recognised as functionally relevant in cytoplasmic capping pathways (Kiss et al. 2015).

### 3.10 *In vitro* Xrn1 susceptibility assay

An *in vitro* Xrn1 susceptibility assay was performed as described before (Islam and Mukherjee 2025). Briefly, A total of 30 μg of cytoplasmic poly(A) selected RNA from different experimental conditions was mixed with capped Renilla luciferase and uncapped GFP spike-in controls at a 1:1000 ratio. The RNA mixture was heat-denatured at 65 °C for 5 min, followed by digestion with 0.5 μL Xrn1 enzyme (NEB, cat. no. M0338S) in the supplied buffer for 1 h at 37 °C. After digestion, RNA was purified by phenol-chloroform extraction followed by ethanol precipitation. RNAs were then reverse transcribed to prepare cDNA using the SuperScript III First-Strand cDNA Synthesis Kit (Invitrogen; cat# 18080051) following the manufacturer’s instructions. Quantitative real-time PCR was performed with specific sets of primers to amplify the target genes. C_t_ values of cCE-targeted transcripts and non-targeted controls were normalised to the capped Renilla luciferase C_t_ obtained for each condition. Loss of 5′ ends was calculated as the difference in normalised C_t_ values before and after Xrn1 digestion. Changes in Xrn1 susceptibility (ΔX) were expressed as the ratio ΔX_K294A_ /ΔX_Control_, where ΔX_Control_ represents relative 5’ end loss in control cells, and ΔX_K294A_ represents the corresponding loss in K294A-expressing cells. The plot showed the relative change in Xrn1 susceptibility between K294A-expressing and control cells.

### 3.11 RNA sequencing and differential gene expression

Cytoplasmic RNA isolated from control and 24 hrs CoCl_2_-treated U2OS cells was subjected to RNA sequencing on the Illumina NovaSeq 6000 platform. RNA integrity was assessed using the TapeStation 4150 (Agilent Technologies) with HS RNA ScreenTape. After quality control, sequencing libraries were prepared using the TruSeq Stranded Total RNA Kit (Illumina; cat# 15032611). Libraries were quantified using a Qubit 4.0 fluorometer (Thermo Fisher Scientific) with the DNA HS Assay Kit (Invitrogen; cat# Q32851) and subsequently sequenced. Read quality was evaluated using FastQC v0.11.9 with default parameters. Adapter trimming was performed using Trimmomatic v0.35, and high-quality reads were aligned to the STAR-indexed human reference genome (GRCh37) using STAR v2.7.9a with default settings.

Differential gene expression analysis was performed using the DESeq2 package in R. Genes with a p-value ≤ 0.05 and log2FC ±1were considered significantly up- or downregulated. Volcano plots were generated using EnhancedVolcano. Genes were grouped based on expression patterns, and functional enrichment analyses (GO, KEGG, Reactome, and Disease Ontology) were performed using AllEnricher.

### 3.12 Bioinformatic analysis for HIF1α binding at *KCTD16,* and *RORA* promoters

Putative HIF1α binding sites on the indicated genes were predicted using the JASPAR online database (http://www.jaspar.genereg.net).

### 3.13 Chromatin immune precipitation (ChIP) assay

HIF1α binding to the KCTD16 and RORA promoters was validated by in vitro chromatin immunoprecipitation (ChIP) using the SimpleChIP Enzymatic Chromatin IP Kit (Cell Signalling Technology; cat# 9003), following a previously standardised protocol (Islam and Mukherjee 2025). Briefly, the CoCl2 treated or untreated cells were cross-linked with 1% formaldehyde for 10 min at room temperature and quenched with glycine. Fixed cells were harvested in ice-cold DPBS supplemented with 1 mM PMSF. Harvested cells were lysed in a lysis buffer (1× buffer A, 0.5 mM DTT, 1× Protease Inhibitor Cocktail, 1 mM PMSF) and nuclei were pelleted down by centrifuging at 2000*g* for 5 min at 4 °C. The nuclear pellet was resuspended in buffer B with 0.5 mM DTT and digested with micrococcal nuclease for 20 min at 37 °C. Following centrifugation, nuclei were resuspended in 1× ChIP buffer and subjected to sonication to obtain fragmented chromatin. After clarification, ∼ 10 μg chromatin was immunoprecipitated overnight at 4 °C with anti-HIF1α antibody or normal rabbit IgG as a negative control. Immune complexes were captured using protein G magnetic beads, washed, and de-crosslinked, and DNA was purified using spin columns. HIF1α binding was analysed by PCR and quantified by qPCR using immunoprecipitated and input DNA. Enrichment was calculated using the % input method, with IgG pull-down serving as the control. Primer sequences are listed in Table 3.

**Table 3:**
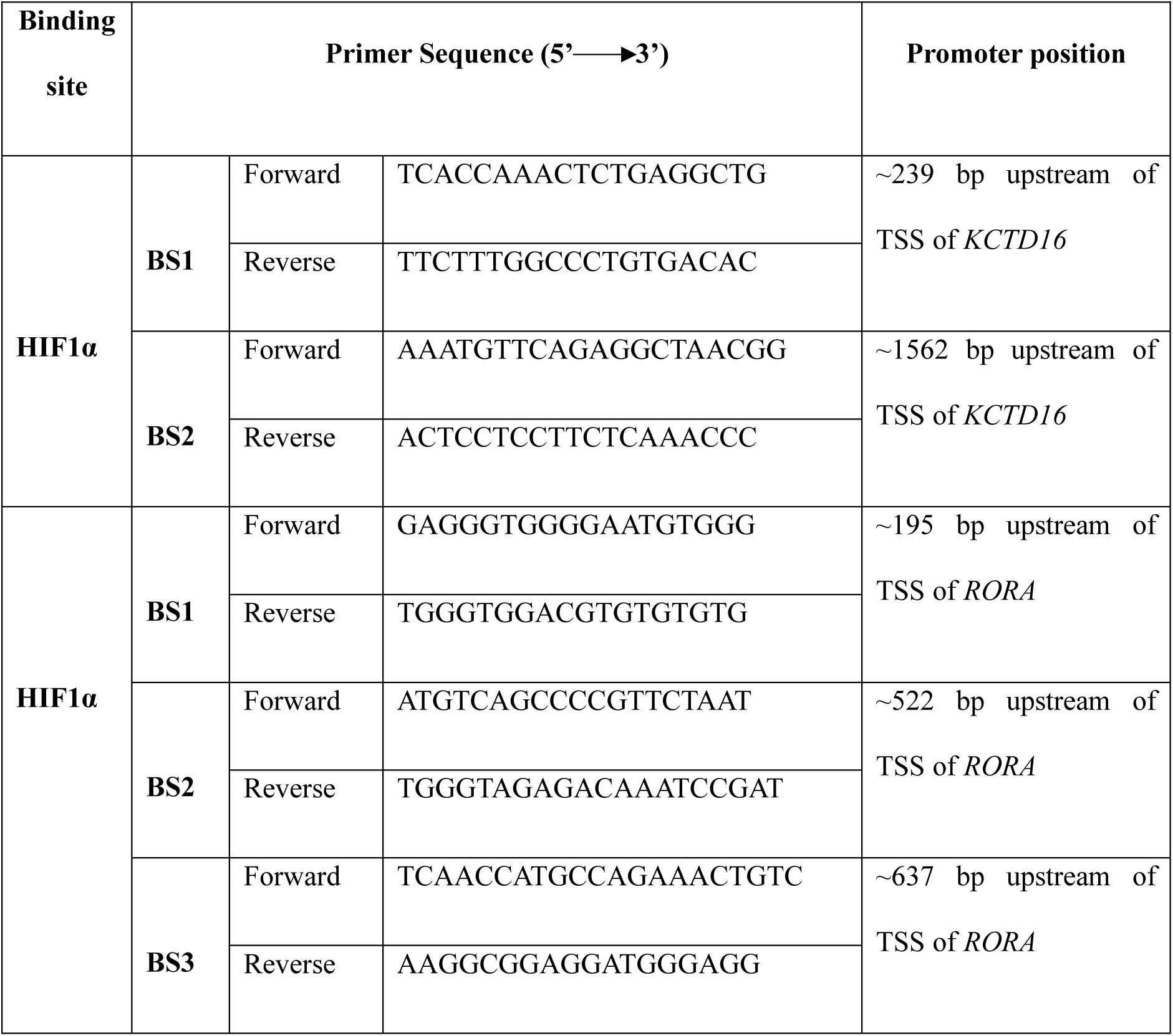
List of ChIP primers used in the study.

### 3.14 Cell cycle analysis by flow cytometry

To assess cell cycle status under CoCl_2_-induced hypoxia, flow cytometry-based cell cycle analysis was performed in U2OS and MG63 cells. Briefly, ∼1 × 10⁶ control or 24 h CoCl_2_-treated cells were fixed in ice-cold 70% ethanol for 30 min at 4 °C, with ethanol added dropwise under gentle vortexing to minimise cell clumping. Fixed cells were washed three times with ice-cold 1× DPBS by centrifugation at 800*g* for 5 min at 4 °C. Cells were then stained with propidium iodide (PI) (Sigma-Aldrich; cat# P4864) at a final concentration of 50 µg/mL, along with RNase A (Invitrogen; cat# 12091021) to remove RNA, and incubated for 30 min at room temperature. A total of 5,000 events per sample were acquired using the S3e Cell Sorter (Bio-Rad), and data were analysed using FCS Express 7 software.

### 3.15 Brdu cell proliferation assay

BrdU cell proliferation assay was performed to analyse the proliferative potential of CoCl_2_ induced hypoxic osteosarcoma cells. Briefly, 70-80% confluent cells were treated with CoCl_2_ or left untreated for 24 hrs, with 10 µM BrdU added during the final 4 hrs of incubation. To evaluate the role of *RORA* and *KCTD16*, cells were treated with 10 µM BrdU for the last 4 hrs, at 48 hrs post-transfection with either overexpression constructs or gene-specific siRNAs. Cells were then washed with 1× DPBS, fixed in chilled methanol for 10 min at -20 °C, and washed three times with DPBS. DNA was denatured with 2 N HCl for 30 min at room temperature, followed by washing and blocking in blocking buffer (5% FBS, 0.2 M glycine, 0.1% Triton X-100 in 1× PBS) for 30 min. Cells were incubated overnight at 4 °C with anti-BrdU antibody (1:200; Abcam; cat# ab6326), washed, and then incubated with Alexa Fluor 350 goat anti-rat IgG (1:200). Cells were counterstained with propidium iodide and treated with RNase A for 1 h at room temperature, washed, and mounted using mounting medium (Invitrogen; cat# S36967). Images were acquired using a Nikon fluorescence microscope with a 20X objective.

### 3.16 Colony formation assay

The colonogenic potential of CoCl_2_ induced hypoxic osteosarcoma cells was examined by performing a conventional colony formation assay. Briefly, Osteosarcoma cells (U2OS or MG63) subjected to the indicated treatments were seeded into 6-well plates or 35-mm dishes at a density of 2 x 10³ cells per well and maintained in complete growth medium in a CO₂ incubator for 10-14 days, with medium changes every 72 hrs. Once visible colonies formed, cultures were washed with 1× PBS, fixed in chilled methanol for 10 min at -20 °C, and stained with 0.05% crystal violet (SRL; cat# 28376) for 30 min at room temperature. Excess stain was removed by washing with tap water, and plates were scanned using the Odyssey CLx Imaging System (LI-COR). Colonies were manually counted.

### 3.17 Wound healing assay

The influence on CoCl_2_ induced hypoxia on the migration of osteosarcoma cells was assessed by performing a wound healing assay. Briefly, A confluent monolayer of U2OS or MG63 cells was scratched using a sterile plastic tip to create a cell-free area and then treated with 200 µM CoCl_2_ or left untreated for 24 hrs at 37 °C in a humidified CO_2_ incubator. Brightfield images were captured using a Leica microscope, and the percentage of wound closure was calculated using the following equation:

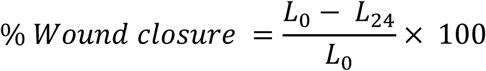

Where *L*_0_ is the initial wound width, and *L*_24_ is the wound width after 24 hrs.

### 3.18 Cell invasion assay

To assess the invasive potential of osteosarcoma cells under CoCl_2_-induced hypoxia, a two-chamber Matrigel invasion assay was performed. Briefly, treated cells were seeded in serum-free medium onto Matrigel-coated inserts (upper chamber), while the lower chamber contained medium supplemented with 10% FBS as a chemoattractant. The setup was incubated at 37 °C in a humidified CO_2_ incubator for 24 hrs. After incubation, non-invaded cells were removed by gentle cotton swabbing, and invaded cells on the underside of the membrane were fixed in chilled methanol and stained with 0.05% crystal violet for 30 min at room temperature. Stained cells were imaged using a bright-field microscope, and invaded cells were counted manually.

### 3.19 Statistical analysis

All statistical analyses were performed using GraphPad Prism 9 (GraphPad Software, Inc.). Two-tailed Student’s t-tests were used for comparisons between two groups, and one-way ANOVA was applied for multiple-group comparisons. Data are presented as mean ± SD from at least three independent biological replicates (n ≥ 3), and a p-value ≤ 0.05 was considered statistically significant.

## SUPPLEMENTAL MATERIAL

Supplemental material is available for this article.

## ACKNOWLEDGMENTS

This work was supported by extramural CSIR-ASPIRE grant from Council of Scientific and Industrial Research, Government of India to Dr. Chandrama Mukherjee. The authors also thank UGC, Government of India for a UGC-SRF fellowship to Mr. Safirul Islam and the Department of Biotechnology, India for a Ramlingaswami Re-entry fellowship to Dr. Chandrama Mukherjee. The authors thank Dr. Shubhra Majumder, Presidency University for providing access to Zeiss Microscope and Dr. Somsubhra Nath, Presidency University for providing reagents for Cell invasion assay and anti-rabbit myc antibody, the Institute of Health Sciences, Presidency University for the departmental instrument facility and Presidency University for necessary infrastructural support.

## Author contributions: CRediT

**Safirul Islam:** Conceptualization, Methodology, Validation, Investigation, data curation, Formal Analysis, Visualization, software, Writing-original draft, Writing - Review & Editing; **Utpal Bakshi:** Methodology, data curation, Formal Analysis, Visualization, software, **Chandrama Mukherjee:** Conceptualization, Resources, Writing - Original Draft, Writing - Review & Editing, Supervision, Project administration, Funding acquisition.

